# Interrogating contrastive learning embeddings for structure-based virtual screening: a case study on DrugCLIP

**DOI:** 10.64898/2026.09.16.752029

**Authors:** Javier S. Utgés, David T. Jones, Christine Orengo

## Abstract

Virtual screening has become central to early-stage drug discovery, and structure-based approaches have recently been reframed as a retrieval problem through contrastive learning methods such as DrugCLIP, which project protein pockets and ligands into a shared embedding space. However, what these abstract representations encode and how they relate to conventional notions of structural and chemical similarity remain unclear. Here we present a systematic dissection of DrugCLIP’s latent space. We show that its pocket embeddings, despite not being explicitly trained for this task, achieve state-of-the-art performance in pocket similarity search while running over 10,000 times faster than existing structural descriptors. We further demonstrate that this embedding space is structurally coherent and robust to conformational variation. Ligand embeddings, by contrast, capture a pocket-aware notion of chemical similarity that extends beyond conventional fingerprint-based measures. Using a rigorous de-leakage benchmark, we show that DrugCLIP generalises beyond its fine-tuning set, retrieving the crystallographic ligand within the top 1% of a library of 48,999 compounds for 56.5% of 223 pockets with low protein sequence identity (SI ≤ 30%) and limited ligand chemical similarity (TS ≤ 0.7) to the fine-tuning set. Nevertheless, performance declines under increasingly realistic screening conditions, primarily due to differences in sidechain orientation across apo, holo and AlphaFold-predicted structures, and mismatches in pocket residue composition when using predicted pockets. These findings delineate the practical limits of DrugCLIP’s applicability, highlight accurate pocket identification and representation as key factors for improving performance, and provide a transferable framework for interpreting the latent spaces of related contrastive pocket-ligand models. Together, these results offer insight to guide the refinement of existing methods and the development of future contrastive screening approaches.

**Scientific contributions:** This work provides the first systematic dissection of the representations learned by DrugCLIP – or any contrastive learning method for virtual screening – establishing that its training objective yields pocket embeddings that not only distinguish binders from non-binders but also encode sufficient structural information to achieve state-of-the-art performance in pocket similarity search, while operating over four orders of magnitude faster than existing descriptors. We further show that DrugCLIP’s embedding space is structurally coherent and robust to conformational variation, with pocket embedding similarity closely reflecting geometric resemblance and ligand embedding similarity capturing a pocket-aware notion of chemical similarity that extends beyond conventional molecular fingerprints. Through a rigorous de-leakage benchmark, we demonstrate that DrugCLIP generalises beyond the protein-ligand complexes encountered during fine-tuning, rather than merely memorising these associations. We also quantify its performance degradation across increasingly realistic screening scenarios – including alternative apo and holo structures, AlphaFold models, and predicted pockets – attributing this degradation to sidechain orientation and differences in residue membership, respectively. These findings clarify the practical boundaries of DrugCLIP’s applicability, identify pocket prediction accuracy as a key element for improving performance when experimental ligand data is not available, and offer a transferable analytical framework for interpreting the latent spaces of related contrastive pocket-ligand encoders such as DrugHash, BindCLIP or ConGLUDe.

**TOC Graphic:** 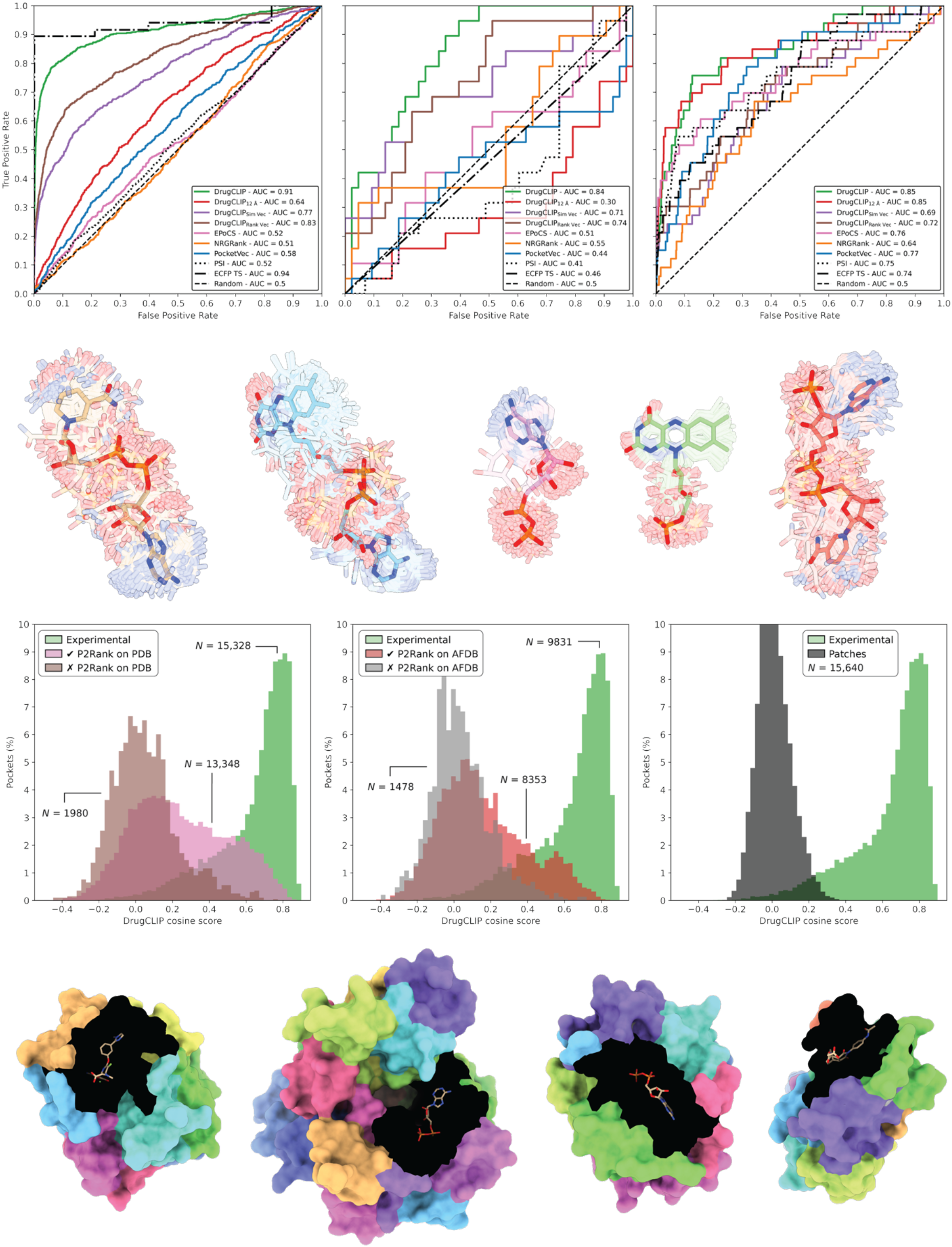

## Introduction

Drug discovery is a complex and resource-intensive process that can take up to 15 years and cost more than $1 billion [1]. These substantial financial and time commitments are largely driven by the high rate of failure, or attrition, experienced by drug candidates throughout development. Computational approaches, such as virtual screening, have therefore become valuable tools for mitigating these challenges. Over the past few decades, virtual screening has contributed to reducing the time and cost associated with early-stage drug discovery by computationally prioritising candidate molecules before undertaking costly experimental validation [2–4]. By narrowing large chemical libraries down to a tractable set of promising compounds, virtual screening enables researchers to focus experimental efforts on candidates with a greater likelihood of success, rather than exhaustively testing millions of molecules.

Historically, the limited availability of experimentally determined three-dimensional (3D) protein structures has been a major bottleneck for structure-based virtual screening, restricting its application to a relatively small subset of well-characterised targets [5, 6]. However, recent advances in protein structure prediction – particularly those driven by deep learning methods such as AlphaFold [7, 8] – have dramatically expanded the coverage of protein targets with reliable structural models available [9, 10]. This progress has broadened the applicability of structure-based virtual screening to previously understudied or experimentally challenging targets, creating new opportunities for drug discovery across a wider range of disease-relevant proteins [11–15].

This unprecedented expansion in available structural data, however, has outpaced the capabilities of many established virtual screening methods. Molecular docking tools [16–19] remain fundamentally constrained by the need to sample individually and score each protein-ligand pair, a process that quickly becomes computationally prohibitive when scaled to proteome-wide target sets and billions of compounds [20]. Deep learning-based scoring methods [21, 22], including approaches based on graph neural networks [23, 24], have emerged as potentially faster alternatives, but their ability to generalise reliably to novel targets and unexplored chemical space remains a significant challenge [25–27]. As a result, a growing number of AI-based methods have recently been proposed to bridge the widening gap between structural data availability and virtual screening capacity [28–30].

Contrastive language-image pretraining (CLIP) was originally developed to align images and their textual descriptions within a shared embedding space, bringing representations of matching pairs closer together while separating those of non-matching pairs [31]. CLIP-inspired approaches have since been successfully applied to diverse tasks such as enzyme function prediction [32], homologue detection [33] and structure classification [34], and more recently to virtual screening, reformulating it as a retrieval rather than pairwise scoring problem [35]. These methods employ two separate encoders – one for the protein pocket and another for the candidate molecule – to project their respective representations into a shared embedding space. A contrastive learning objective encourages known binding pairs to have similar representations, while separating them from non-binding pairs. Crucially, ligand embeddings can be precomputed and reused across queries, allowing new targets to be screened against libraries containing millions or billions of compounds through efficient similarity searches, rather than repeated evaluations of individual protein-ligand pairs. DrugCLIP [36] pioneered this approach to virtual screening, and several methods have adopted or extended this paradigm since, including LigUnity [37], HypSeek [38] and SPRINT [39].

Despite these advances, it remains unclear what these abstract representations capture exactly, and how they relate to conventional measures of structural and chemical similarity. Previous studies have explored these questions only indirectly – for instance, by testing robustness to apo, holo, and AlphaFold-predicted structures [35, 36, 40], comparing performance under different pocket prediction methods [41], or examining the physicochemical properties of retrieved molecules against those identified through fingerprint-based searches [42]. However, these analyses did not directly characterise the information encoded within the pocket and ligand embeddings themselves. Here, we address this gap through a systematic dissection of DrugCLIP’s latent space. We demonstrate that its pocket embeddings achieve state-of-the-art performance in pocket similarity search across the ProSPECCTs benchmarks, despite not having been explicitly trained for this task, while operating several orders of magnitude faster than existing pocket descriptors. We further show that DrugCLIP’s representations capture meaningful structural relationships, exhibit robustness to conformational variation, and support generalisation to proteins and ligands beyond its fine-tuning set. Finally, we quantify the deterioration in screening performance under increasingly realistic conditions, highlighting sidechain orientation and differences in pocket residue membership as key contributing factors. Although our analysis focuses on DrugCLIP, the proposed framework is, in principle, applicable to other methods that reformulate virtual screening using independently encoded pocket and ligand representations, including ConGLUDe [41], DrugHash [43] and BindCLIP [44].

## Results

### Pocket matching benchmark

Figure 1, Table 1 and Table S1 summarise the results of the pocket matching benchmark on the ProSPECCTs [45] suite of data sets, which evaluates the ability of different methods to discriminate between similar and dissimilar pocket pairs. We compare DrugCLIP and three novel variants introduced in this work against recently developed methods, including EPoCS [46], NRGRank [47] and PocketVec [48], alongside simple baselines for reference. Notably, the “PSI” baseline, which classifies pocket pairs based on pocket sequence identity, achieve perfect discrimination, area under the curve (AUC) = 1.0, on data sets D1, D1.2, D2, D3, and D4, while the “Sequence” baseline, based on receptor protein sequence identity, achieves the same only on D2. These results reflect the construction of these data sets, in which positively labelled pocket pairs originate from the same protein. Consequently, high AUROC values on D1, D1.2 and D2 should be interpreted cautiously, as they may primarily reflect the recognition of identical or closely related protein pockets rather than generalisable pocket similarity across diverse proteins. Similarly, the “Ligand Code” and “ECFP4 TS” baselines achieve AUROC ≈ 0.95 on D5 and D5.2, where pockets from different proteins binding identical or chemically similar ligands are labelled as positive pairs. Although these baselines outperform more sophisticated methods on certain data sets, their performance varies across evaluation scenarios, and they are insufficient on their own to provide a general quantitative measure of pocket similarity across experimentally determined protein-ligand complexes, let alone estimated or predicted ligand binding pockets.

**Figure 1.**
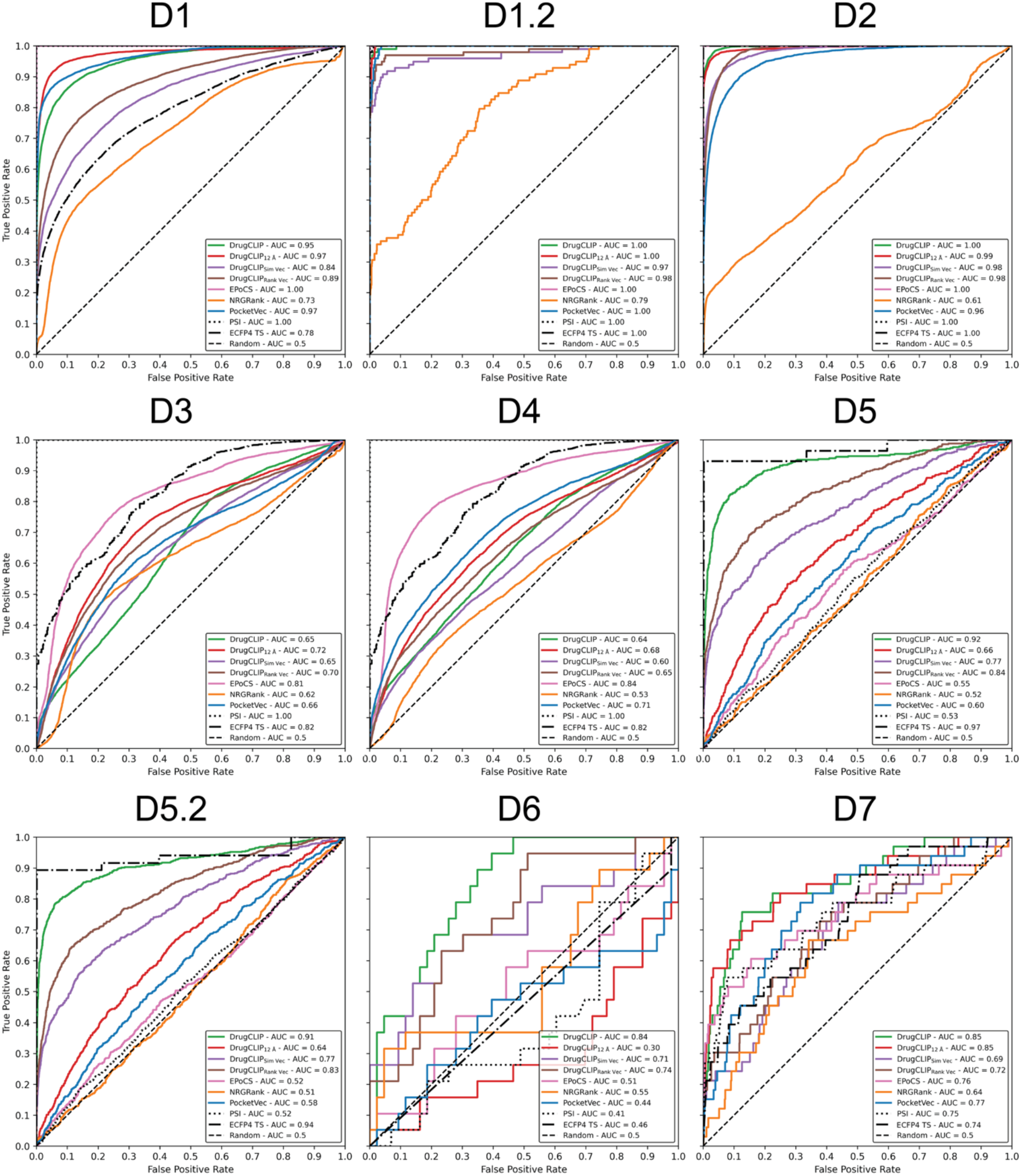
ProSPECCTs pocket matching benchmark. Summary of the pocket matching performance across the nine ProSPECCTs data sets evaluated in this study. See Table 4 for the number of pockets and pairwise comparisons, including positive and negative pairs, in each data set. D6.2 was excluded because its cofactor atoms are incompatible with DrugCLIP’s pocket encoder. ROC curves and their corresponding AUCs are shown for each data set. DrugCLIP, EPoCS and PocketVec were evaluated using their default configurations, while NRGRank was used to screen the same 128 lead-like molecules employed by PocketVec, generating pseudo-docking-derived pocket descriptors following an analogous approach. Three additional DrugCLIP variants were evaluated: DrugCLIP_12 Å_, which defines pockets using a 12 Å threshold instead of the default 6 Å; DrugCLIP_Sim Vec_, which uses DrugCLIP’s screening scores against the same 128 PocketVec molecules to generate pocket-ligand similarity vectors; and DrugCLIP_Rank Vec_, which instead represents pockets through the rankings of these molecules based on their screening scores. Two baselines are also included for reference, classifying pocket pairs based on pocket sequence identity (PSI) or the Tanimoto similarity between bound ligand ECFP4 fingerprints (ECFP4 TS).

**Table 1.** ProSPECCTs pocket matching benchmark. Summary of the pocket matching performance across the nine ProSPECCTs sets evaluated in this study. All reported values are AUROC. Two additional naïve binary baselines are included for reference, which classify pocket pairs based on identical receptor protein sequences, determined by sequence hashes, or 100% sequence identity (“Sequence”), or identical bound ligand codes (“Ligand Code”).

|  | **D1** | **D1.2** | **D2** | **D3** | **D4** | **D5** | **D5.2** | **D6** | **D7** |
|---|---|---|---|---|---|---|---|---|---|
| DrugCLIP | 0.95 | **1.00** | **1.00** | 0.65 | 0.64 | **0.92** | **0.91** | **0.84** | **0.85** |
| DrugCLIP<sub>12 Å</sub> | 0.97 | **1.00** | 0.99 | 0.72 | 0.68 | 0.66 | 0.64 | 0.30 | **0.85** |
| DrugCLIP<sub>Sim Vec</sub> | 0.84 | 0.97 | 0.98 | 0.65 | 0.60 | 0.77 | 0.77 | 0.71 | 0.69 |
| DrugCLIP<sub>Rank Vec</sub> | 0.89 | 0.98 | 0.98 | 0.70 | 0.65 | 0.84 | 0.83 | 0.74 | 0.72 |
| EPoCS | **1.00** | **1.00** | **1.00** | **0.81** | **0.84** | 0.55 | 0.52 | 0.51 | 0.76 |
| NRGRank | 0.73 | 0.79 | 0.61 | 0.62 | 0.53 | 0.52 | 0.51 | 0.55 | 0.64 |
| PocketVec | 0.97 | **1.00** | 0.96 | 0.66 | 0.71 | 0.60 | 0.58 | 0.44 | 0.77 |
| Sequence | 0.66 | 0.87 | **1.00** | 0.66 | 0.66 | 0.50 | 0.50 | 0.50 | 0.50 |
| Ligand Code | 0.50 | 0.50 | 0.99 | 0.50 | 0.50 | **0.97** | **0.95** | 0.47 | 0.60 |
| PSI | **1.00** | **1.00** | **1.00** | **1.00** | **1.00** | 0.53 | 0.52 | 0.41 | 0.75 |
| ECFP4 TS | 0.78 | **1.00** | **1.00** | 0.82 | 0.82 | **0.97** | 0.94 | 0.46 | 0.74 |

We consider D5, D5.2, D6 and D7 particularly relevant within the ProSPECCTs suite, as they address a central application of pocket similarity search: identifying binding sites capable of recognising similar ligands across structurally and evolutionarily diverse proteins. We therefore argue that strong performance on these data sets is more informative for this application than performance on D1 and D2, which primarily assess robustness to structural variation, or the synthetic D3 and D4 sets, which evaluate sensitivity to artificially introduced residue substitutions.

DrugCLIP pocket embeddings demonstrate strong discriminatory power across seven of the nine ProSPECCTs sets (AUROC ≥ 0.80), including D1, D1.2 and D2, where DrugCLIP, EPoCS and PocketVec all achieve an AUROC ≥ 0.95. EPoCS achieves perfect discrimination (AUROC = 1.0) on these three sets, consistent with its reliance on sequence-based representations and the fact that positively labelled pocket pairs originate from the same protein. EPoCS also outperforms all other methods on D3 and D4, suggesting that the artificially introduced residue substitutions in these data sets have a greater impact on the sequence embeddings employed by EPoCS than on the representations learned by DrugCLIP or the docking-derived rank vectors used by NRGRank and PocketVec.

DrugCLIP consistently outperforms EPoCS, NRGRank and PocketVec, on D5, D5.2, D6 and D7. Its strong performance on D5 and D5.2 indicates that its learned representations capture sufficient detail of the binding pocket environment to recognise similarities between pockets from evolutionarily divergent proteins that bind similar ligands [49]. Performance on D6 further suggests that DrugCLIP captures aspects of ligand binding modes beyond the recognition of pockets accommodating similar ligands [50]. Finally, its performance on D7 supports the generalisability of DrugCLIP’s pocket representations across a diverse collection of similar binding sites reported in the literature [51].

The alternative DrugCLIP variants evaluated in this benchmark generally underperform relative to the default approach, which calculates similarity directly between pocket embeddings defined using residues within 6 Å of the ligand. Expanding the pocket definition to include residues within 12 Å substantially reduces discriminatory performance, with AUROC decreases (Δ_AUC_) of up to 0.27 on D5.2 and 0.54 on D6. This suggests that incorporating the broader structural environment, including residues that do not directly interact with the ligand, introduces potentially confounding information that reduces the ability of DrugCLIP embeddings to distinguish between similar and dissimilar pockets. Among the “Sim Vec” and “Rank Vec” variants, the latter outperforms the former on eight of the nine sets, indicating that ligand rankings derived from DrugCLIP-based screening, analogous to the approach employed by PocketVec, provide greater discriminatory power than the raw screening scores.

NRGRank underperforms relative to the other methods on eight of the nine ProSPECCTs data sets, suggesting that it is less effective at generating pocket descriptors using the same approach employed by PocketVec with rDock [52], at least when using the same library of 128 lead-like compounds [48]. This may be attributable to NRGRank’s use of approximate, knowledge-based scoring functions, rather than the more physically grounded energy calculations employed by conventional docking tools. However, this simplification also contributes to NRGRank being approximately 100-fold faster than traditional docking methods [47]. Given this computational efficiency, screening a larger and more chemically diverse compound library could potentially improve its discriminatory performance, bringing it closer to, or even beyond, that of PocketVec.

Whilst EPoCS performs well on D1, D1.2, D2, D3 and D4, its performance deteriorates substantially on D5, D5.2 and D6, highlighting its limited ability to recognise similar binding pockets across structurally diverse proteins. This is consistent with its reliance on sequence-based ESM-2 embeddings, which do not explicitly account for ligand binding properties. PocketVec, in contrast, approaches pocket characterisation from a different perspective, deriving its descriptors from ligand docking calculations rather than directly encoding the structural environment of the binding site, as DrugCLIP does. Furthermore, while PocketVec requires approximately one hour to characterise a single pocket [48], DrugCLIP generates pocket embeddings in less than a second, representing a speedup of over 10,000-fold (Table S4). This substantial computational advantage makes DrugCLIP particularly well suited for large-scale pocket characterisation, comparison and similarity searches.

Tables S2 and S3 extend this evaluation to include the sixteen methods [53–66] assessed by Ehrt *et al.* [45] in 2018, alongside five more recently developed approaches: DeeplyTough [67], Site2Vec [68], BindSiteS-CNN [69], SiteMine [70] and SiteContext [71]. The inclusion of these 21 additional methods reinforces our findings, demonstrating that DrugCLIP not only outperforms recent docking-derived ligand response-profile methods, such as PocketVec and NRGRank, but also achieves superior overall performance across the ProSPECCTs suite compared with deep learning-based approaches (EPoCS, DeeplyTough and BindSiteS-CNN), earlier pharmacophore- and fingerprint-based methods (FuzCav, KRIPO and TIFP), and alignment-based tools (SMAP, Shaper and SiteMine).

### DrugCLIP representations’ analysis

To better understand the abstract representations, or embeddings, generated by DrugCLIP’s pocket and ligand encoders, we conducted a series of analyses using four data sets of protein-ligand complexes widely employed for training and evaluating ligand binding site prediction methods [72–75]: COACH420 [76], HOLO4K [77], scPDB [78] and PDBBind [79]. Together, these data sets encompass a diverse range of binding sites and molecules, including naturally occurring ligands, pharmacologically relevant complexes with experimentally measured binding affinity, and single- and multi-chain protein structures. Further details are provided in the Methods section.

DrugCLIP’s molecule encoder was trained using a random conformer sampling strategy, better reflecting realistic virtual screening scenarios in which the bound conformation of a ligand is unknown. This training approach is expected to improve the model’s robustness to conformational variability and enhance its generalisation capabilities [36]. To assess the sensitivity of DrugCLIP’s ligand representations to conformational differences, we calculated cosine similarities between embeddings of 11,066 conformers from 1461 distinct ligands represented by multiple conformations across the four protein-ligand data sets. Figure 2 A-E illustrates the five most frequently occurring ligands: nicotinamide-adenine-dinucleotide (NAD), flavin adenine dinucleotide (FAD), adenosine-5’-diphosphate (ADP), flavin mononucleotide (FMN) and nicotinamide-adenine-dinucleotide phosphate (NAP), represented by 604, 548, 463, 320 and 263 conformers, respectively. Figure 2 F shows the mean pairwise cosine similarity between conformer embeddings of each ligand, which exceeds 0.9 for all molecules, demonstrating that DrugCLIP’s molecule encoder generates highly consistent representations across different conformations of the same molecule.

**Figure 2.**
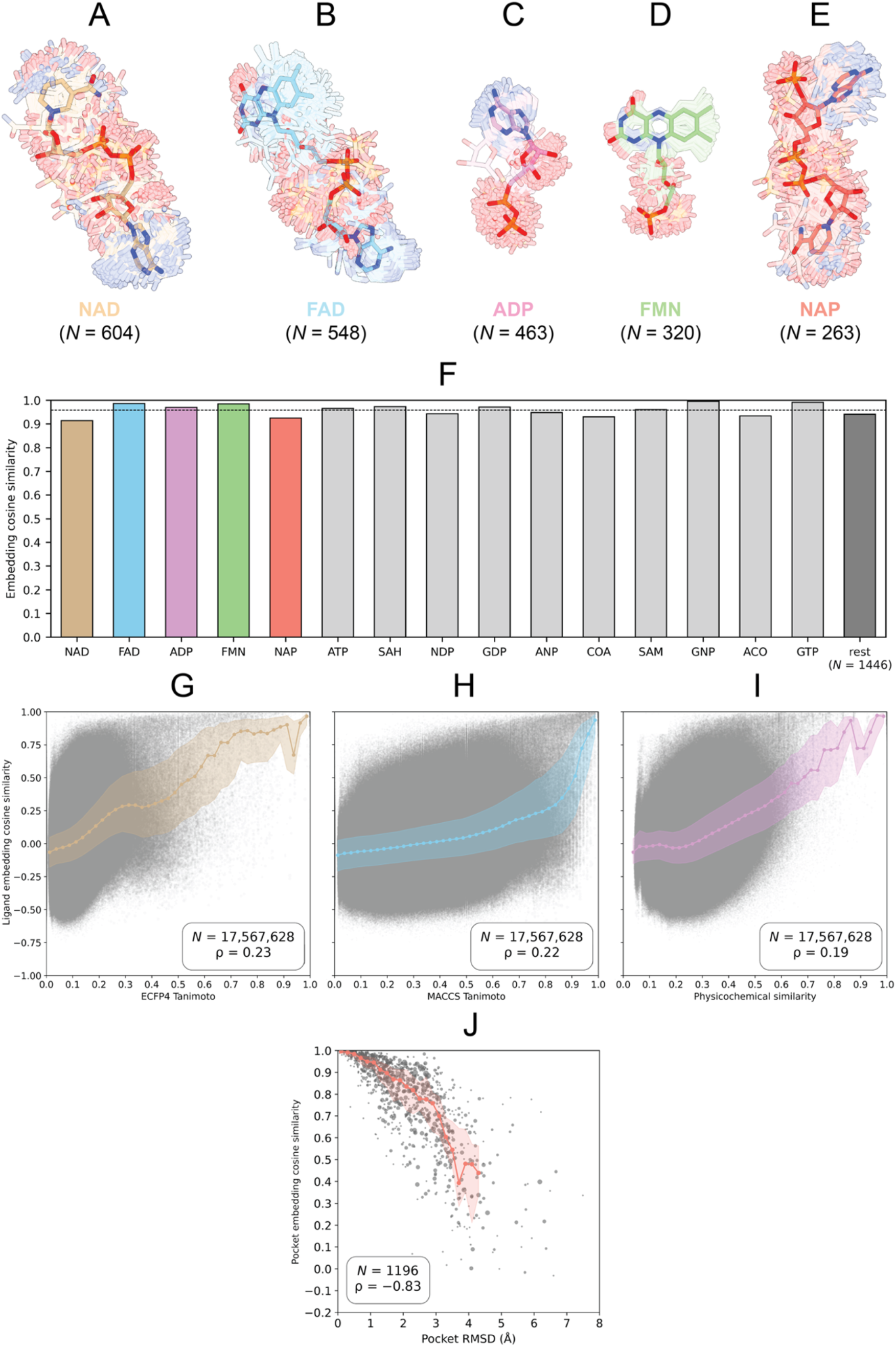
DrugCLIP representations analysis. Characterisation of the pocket and ligand embeddings learned by DrugCLIP across four protein-ligand data sets: COACH420, HOLO4K, scPDB and PDBBind. **(A-E)** Structural superposition of the five most frequently occurring ligands across these data sets. **(A)** Nicotinamide-adenine-dinucleotide (NAD) – *N* = 604; **(B)** Flavin-adenine dinucleotide (FAD) – *N* = 548; **(C)** Adenosine-5’-diphosphate (ADP) – *N* = 463; **(D)** Flavin mononucleotide (FMN) – *N* = 320; **(E)** Nicotinamide-adenine-dinucleotide phosphate (NAP) – *N* = 263; **(F)** Mean pairwise embedding cosine similarity between conformers of the same ligand – *N* = 1461. The overall mean similarity (0.96) is indicated by a black dashed horizontal line. The five ligands shown in **A-E** are highlighted in tan, sky blue, plum, light sea green and salmon, respectively; **(G-I)** Relationships between ligand embedding cosine similarity and chemical similarity, measured using **(G)** ECFP4 Tanimoto similarity (TS) – *N* = 17,567,628, ρ = 0.23; **(H)** MACCS keys TS – *N* = 17,567,628, ρ = 0.22; and **(I)** physicochemical similarity derived from a 10-descriptor profile comprising molecular weight, logP, TPSA, hydrogen-bond donor and acceptor counts, rotatable bond count, aromatic ring count, Csp^3^ fraction, ring count and heavy atom count). Physicochemical similarity was calculated as 1/(1 + Euclidean distance) – *N* = 17,567,628, ρ = 0.19. Each data point represents the mean embedding similarity across all instances of a given ligand pair; **(J)** Relationship between pocket embedding cosine similarity and structural similarity, measured by sidechain atoms RMSD. Each data point represents the mean embedding similarity and RMSD across all instances of the same-ligand pocket pair comparisons – *N* = 1196, ρ = −0.83. Marker sizes are scaled logarithmically according to the number of pocket-pair comparisons represented by each data point. Only pocket pairs which ligand RMSD ≤ 1 Å were included to ensure reliable structural alignment. Spearman’s rank correlation coefficient (ρ) was used throughout to assess monotonic relationships without assuming linearity.

We next examined the extent to which DrugCLIP’s ligand embeddings capture chemical similarity, as measured by the Tanimoto coefficient (TC) between extended connectivity fingerprints (ECFP4) and MACCS keys, as well as similarity derived from physicochemical properties. Figure 2 G-I illustrates the relationships between these chemical similarity measures and DrugCLIP molecule embedding cosine similarities, calculated from 17,567,628 pairwise comparisons among 5928 unique ligands across the four protein-ligand data sets. Weak positive correlations were observed across all three measures – Spearman’s ρ = 0.23 for ECFP4, ρ = 0.22 for MACCS keys, and ρ = 0.19 for physicochemical similarity (all p Δ 0) – indicating that chemically related molecules tend to have more similar DrugCLIP embeddings, although this relationship is limited regardless of the chemical similarity measure employed. Focusing on ECFP4, most ligand pairs exhibit Tanimoto similarities within [0.0, 0.4], yet span a broad range of cosine similarities [–0.5, 0.75], suggesting that chemically dissimilar molecules can nevertheless occupy nearby regions of DrugCLIP’s latent space. This may reflect the fact that DrugCLIP is not trained to predict chemical similarity directly, but rather to learn representations that favour compatible pocket-ligand pairs. Consequently, chemically dissimilar ligands capable of binding to similar pockets may be positioned close together in the embedding space. DrugCLIP therefore appears to capture aspects of ligand similarity beyond conventional chemical descriptors, potentially reflecting molecular features relevant to compatibility with similar binding environments.

To further investigate the pocket-aware nature of DrugCLIP ligand embeddings, we stratified ligand pairs according to the cosine similarity between the embeddings of their corresponding binding pockets (Figure S1). The correlation between ligand embedding cosine similarity and ECFP4 Tanimoto similarity increased progressively with pocket similarity, from ρ = 0.08 for the most dissimilar pocket pairs – cosine similarity ∈ [−0.75, −0.50] – to ρ = 0.62 for the most similar pairs – cosine similarity ∈ (0.75, 1.00] – (Figure S1 A-G). Furthermore, pocket and ligand embedding similarities were themselves positively correlated – ρ = 0.57 – (Figure S1 H), whereas pocket embedding similarity showed only a weak correlation with ligand ECFP4 Tanimoto similarity – ρ = 0.13 – (Figure S1 I). Together, these findings provide further evidence that DrugCLIP ligand embeddings capture aspects of chemical similarity that are influenced by the characteristics of the corresponding binding environments and are not fully explained by conventional molecular fingerprints.

Figure 2 J explores the association between pocket embedding similarity and structural similarity, measured by sidechain atom root mean square deviation (RMSD), to assess the extent to which DrugCLIP’s pocket representations reflect structural differences. A total of 716,099 pocket pairs, comprising 11,121 unique pockets binding 1479 ligands, were analysed. Pockets binding the same ligand were structurally aligned using the bound molecule as a reference, and pocket RMSD was calculated from the transformed sidechain atom coordinates. To ensure reliable alignment of the pocket atoms interacting with the ligand, the analysis was restricted to pocket pairs with a bound-ligand RMSD ≤ 1 Å, yielding 147,791 pairs (20.6%) comprising 9643 unique pockets (86.7%) and 1196 ligands (80.9%). A strong negative correlation was observed between pocket embedding cosine similarity and RMSD (ρ = –0.83; p Δ 0), with structurally similar pockets (low RMSD) exhibiting high embedding similarity, whereas structurally divergent pockets (high RMSD) generally displayed lower similarity scores. These findings demonstrate that DrugCLIP’s pocket embeddings closely reflect the structural organisation of binding sites, suggesting that pocket geometry and sidechain orientation play important roles in determining pocket-pocket and pocket-ligand compatibility.

### Bound ligand recovery benchmark

Building on the virtual screening evaluation of DrugCLIP by Jia *et al.* [36], which employed the DUD-E [80–82] and LIT-PCBA [83] reference data sets, we assess DrugCLIP on the related task of bound ligand recovery using COACH420, HOLO4K, scPDB and PDBBind. Given a ligand-stripped query pocket from an experimentally determined protein-ligand complex deposited in the PDBe [84, 85], and a library of candidate compounds containing the experimentally observed ligand, DrugCLIP was tasked with recovering the bound molecule by screening the entire library against the query pocket and ranking compounds according to their protein-ligand embedding cosine similarity scores. A library of 48,999 non-ionic CCD [86] ligands containing more than one atom was employed for this purpose. To investigate the extent to which DrugCLIP relies on memorisation of its fine-tuning data and assess its ability to generalise beyond them, we evaluated increasingly challenging subsets of pocket-ligand complexes with progressively reduced overlap with the fine-tuning set.

Figure 3 and Table 2 summarise the results of the bound ligand recovery benchmark. DrugCLIP demonstrated strong performance across the pooled set of 15,533 pockets, achieving an EF_1%_ of 84.8, a top 1 recall of 12.9%, and a median rank of 12. This indicates that the experimentally observed ligand was retrieved within the top 1% of the 48,999-compound library for 84.8% of pockets, ranked first for 12.9%, and placed at rank #12 or better for half of the evaluated pockets (*N* = 7766). However, this performance is partly attributable to overlap with DrugCLIP’s fine-tuning set, as 6210 pockets (40%) correspond to structures included in this set, for which performance reaches an EF_1%_ of 96.5, a top 1 recall of 20.5% and a median rank of 5. As progressively lower protein sequence identity thresholds relative to the BioLiP [87] subset used to fine-tune DrugCLIP are imposed, performance declines, reaching an EF_1%_ of 72.7, a top 1 recall of 5.8% and a median rank of 85 for the 962 pockets sharing ≤ 30% sequence identity with the fine-tuning set. Together, filtering by structural overlap and protein sequence identity results in a 12.1-point reduction in EF_1%_ relative to the pooled set (Figure 3 A-C).

**Figure 3.**
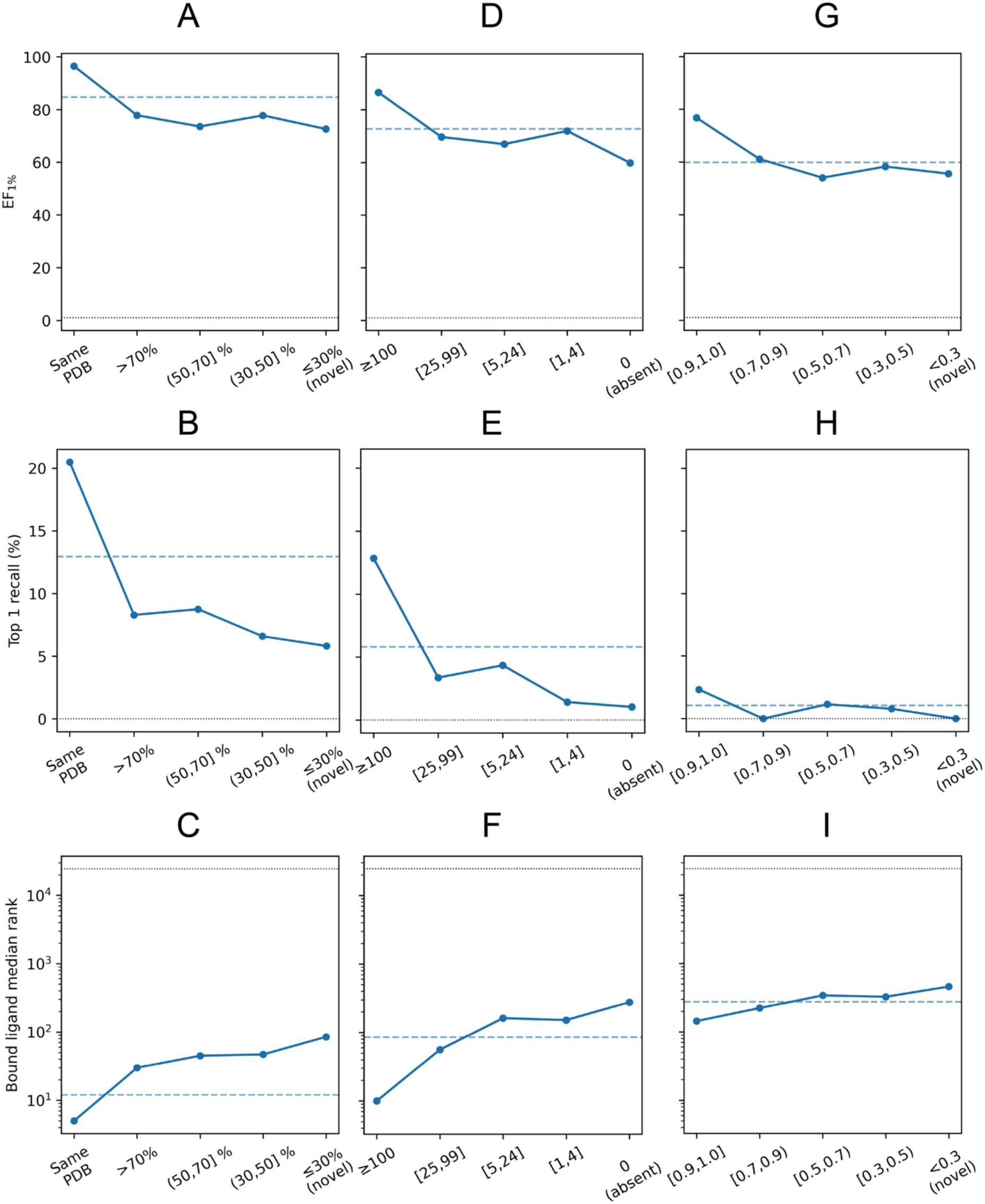
Bound ligand recovery benchmark. Summary of DrugCLIP’s performance on the bound ligand recovery task and its dependence on potential training-test overlap with the BioLiP subset used for fine-tuning. Leakage, memorisation and generalisation were investigated through three sequential and cumulative filtering levels, each comprising several tiers: protein sequence similarity, ligand frequency and ligand chemical similarity. At the protein level, complexes were first assessed for PDB overlap with the fine-tuning set, followed by stratification according to protein sequence identity (SI). At the ligand frequency level, the remaining complexes were stratified by the frequency of their bound ligands in the fine-tuning set, based on CCD identifiers. Finally, complexes containing ligands absent from the fine-tuning set were stratified by their maximum ECFP4 Tanimoto Similarity (TS) to ligands encountered during fine-tuning. **(A-C)** Effect of PDB overlap and protein sequence similarity on **(A)** EF_1%_, **(B)** top 1 recall (%) and **(C)** median bound molecule rank within the CCD library; **(D-F)** Effect of ligand frequency on the same three metrics, respectively, restricted to complexes with protein SI ≤ 30%; **(G-I)** Effect of ligand chemical similarity on the same metrics, restricted to complexes with protein SI ≤ 30% and ligands absent from the fine-tuning set. Blue dashed lines indicate performance on the corresponding starting subsets before stratification: the entire pooled set for **A-C**, complexes with protein SI ≤ 30% for **D-F**, and complexes with protein SI ≤ 30% and containing ligands absent from the fine-tuning set for **G-I**. Black dotted lines indicate random retrieval baselines of EF_1%_ = 1, top 1 recall = 0.002% and median rank = 24,500. The number of pockets evaluated at each de-leakage tier is reported in Table 2.

**Table 2.** Bound ligand recovery benchmark. Summary of DrugCLIP’s performance on the bound ligand recovery task and its dependence on potential training-test overlap with the BioLiP subset used for fine-tuning. Enrichment factor at 1% (EF_1%_), top 1 recall (%), median bound ligand rank and the number of evaluated pockets are reported for each tier across three sequential and cumulative de-leakage levels. Values in brackets indicate 95% confidence intervals for EF_1%_ and top 1 recall, calculated as binomial proportions. The analysis starts with the pooled set of 15,533 pockets from the filtered COACH420, HOLO4K, scPDB and PDBBind data sets [75]. The first level considers PDB overlap and protein sequence identity (SI) relative to the fine-tuning set, with tiers comprising identical PDB codes, SI (%) > 70, SI ∈ (50, 70], SI ∈ (30, 50], SI ≤ 30 (*novel* proteins). The second level examines the frequency of the bound ligand in the fine-tuning set: ≥ 100, 25-99, 5-24, 1-4, and 0 occurrences (*absent* ligands). The final level stratifies the remaining complexes according to the maximum ECFP4 Tanimoto similarity (TS) between their bound ligands and those in the fine-tuning set, using intervals of [0.9, 1.0], [0.7, 0.9), [0.5, 0.7), [0.3, 0.5), and < 0.3 (*chemically dissimilar* ligands). Each successive level is applied exclusively to the subset satisfying the strictest criterion on the preceding level, progressively reducing the number of evaluated pockets from 15,533 to just nine in the most stringent de-leakage tier.

| **Data Bin** | ***N*** | **EF<sub>1%</sub>– 95% CI** | **Recall<sub>top 1</sub> (%) – 95% CI** | **Median Rank** |
|---|---|---|---|---|
| Entire pooled dataset | 15,533 | 84.8 [84.2, 85.3] | 12.9 [12.4, 13.5] | 12 |
| Exact PDB match | 6210 | 96.5 [96.0, 96.9] | 20.5 [19.5, 21.5] | 5 |
| % SI > 70 | 6618 | 77.9 [76.8, 78.8] | 8.3 [7.7, 9.0] | 30 |
| % SI ∈ (50, 70] | 788 | 73.6 [70.4, 76.6] | 8.8 [7.0, 10.9] | 45 |
| % SI ∈ (30, 50] | 955 | 77.8 [75.1, 80.3] | 6.6 [5.2, 8.4] | 47 |
| % SI ≤ 30 (*novel*) | 962 | 72.7 [69.8, 75.4] | 5.8 [4.5, 7.5] | 85 |
| # Ligand ≥ 100 | 335 | 86.6 [82.5, 89.8] | 12.8 [9.7, 16.8] | 10 |
| # Ligand ∈ [25, 99] | 89 | 69.7 [59.5, 78.2] | 3.4 [1.2, 9.4] | 56 |
| # Ligand ∈ [5, 24] | 115 | 67.0 [57.9, 74.9] | 4.3 [1.0, 9.8] | 162 |
| # Ligand ∈ [1, 4] | 139 | 71.9 [64.0, 78.7] | 1.4 [0.4, 5.1] | 152 |
| # Ligand = 0 (*absent*) | 284 | 59.9 [54.1, 65.4] | 1.1 [0.2, 6.2] | 277 |
| TS ∈ [0.9, 1.0] | 43 | 76.7 [62.3, 86.8] | 2.3 [0.4, 12.1] | 144 |
| TS ∈ [0.7, 0.9) | 18 | 61.1 [38.6, 79.7] | 0.0 [0.0, 17.6] | 223 |
| TS ∈ [0.5, 0.7) | 87 | 54.0 [43.6, 64.1] | 1.1 [0.2, 6.2] | 343 |
| TS ∈ [0.3, 0.5) | 127 | 58.3 [49.6, 66.5] | 0.8 [0.1, 4.3] | 327 |
| TS < 0.3 (*novel*) | 9 | 55.6 [26.7, 81.1] | 0.0 [0.0, 29.9] | 461 |

Ligand frequency in the fine-tuning set also influences retrieval performance. DrugCLIP performs substantially better on pockets binding ligands observed frequently during fine-tuning – ≥100 occurrences (*N* = 335 pockets) – achieving an EF_1%_ of 86.6, a top 1 recall of 12.8% and a median rank of 10, than on those binding ligands absent from the fine-tuning set – 0 occurrences (*N* = 284) – for which these metrics decrease to 59.9, 1.1% and 277, respectively. Restricting the analysis to ligands absent from the fine-tuning set therefore results in a further 12.8-point reduction in EF_1%_ (Figure 3 D-F). Within this subset of pockets from proteins sharing ≤ 30% sequence identity and binding previously unseen ligands (by CCD code), DrugCLIP performs better when the bound molecules are chemically similar to those encountered during fine-tuning – ECFP4 TS ∈ [0.9, 1.0] – achieving an EF_1%_ of 76.7, a top 1 recall of 2.3% and a median rank = 144 (*N* = 43 pockets). In contrast, performance decreases to an EF_1%_ of 55.6, a top 1 recall of 0.0% and a median rank of 461 for ligands with limited chemical similarity to the fine-tuning set – ECFP4 TS < 0.3. Although only nine pockets satisfy this strictest criterion, the overall trend is consistent across the different TS intervals (Figure 3 G-I).

Considering the 223 pocket-ligand pairs satisfying both ≤ 30% sequence identity and ≤ 0.7 ECFP4 Tanimoto similarity relative to the fine-tuning set, DrugCLIP achieves an EF_1%_ of 56.5 – 95% CI = [49.9, 62.8] – a top 1 recall of 0.9% – 95% CI = [0.3, 3.2] – and a median rank of 347. These values represent a substantial decline from the corresponding results on the full pooled set (84.8, 12.9% and 12, respectively). Nevertheless, DrugCLIP still substantially outperforms random retrieval, achieving a 57-fold improvement in EF_1%_, a 450-fold increase in top 1 recall and a 71-fold reduction in median rank relative to the random baselines (EF_1%_ = 1, top 1 recall = 0.002% and median rank = 24,500).

### Score robustness and degradation analysis

The bound ligand recovery benchmark described above relies on knowledge of both the experimental binding pocket structure and the crystallographic pose of the bound ligand. This contrasts with realistic virtual screening scenarios, where only alternative apo or holo conformations of the target protein may be available, predicted structures may be required, ligand poses are unknown, and binding sites may need to be identified using pocket prediction tools. Here, we investigate how these factors affect DrugCLIP’s performance by examining their impact on pocket-pocket and pocket-ligand embedding cosine similarities.

Figure 4 A compares the distributions of DrugCLIP cosine similarity scores between CCD ligand conformers and their corresponding pockets observed in the reference experimental PDB structures (green) or the equivalent pockets transferred to the AFDB predicted models (purple) across 13,062 unique pockets. The experimental distribution exhibits a pronounced peak around 0.8, whereas the AFDB distribution shows a broader concentration of scores between −0.2 to 0.6. Consistent with this shift, 82.5% of ligands achieve higher scores against their experimental pockets than against their AFDB counterparts, with a median score difference of −0.09 (AFDB relative to PDB). Figure 4 B examines the relationship between structural similarity, measured by sidechain (SC) root mean square deviation (RMSD), and embedding (cosine) similarity between experimental (canonical) and AFDB pockets. A strong negative Spearman correlation (ρ = –0.67) indicates that pockets with greater structural deviations tend to be more distant in DrugCLIP’s embedding space.

**Figure 4.**
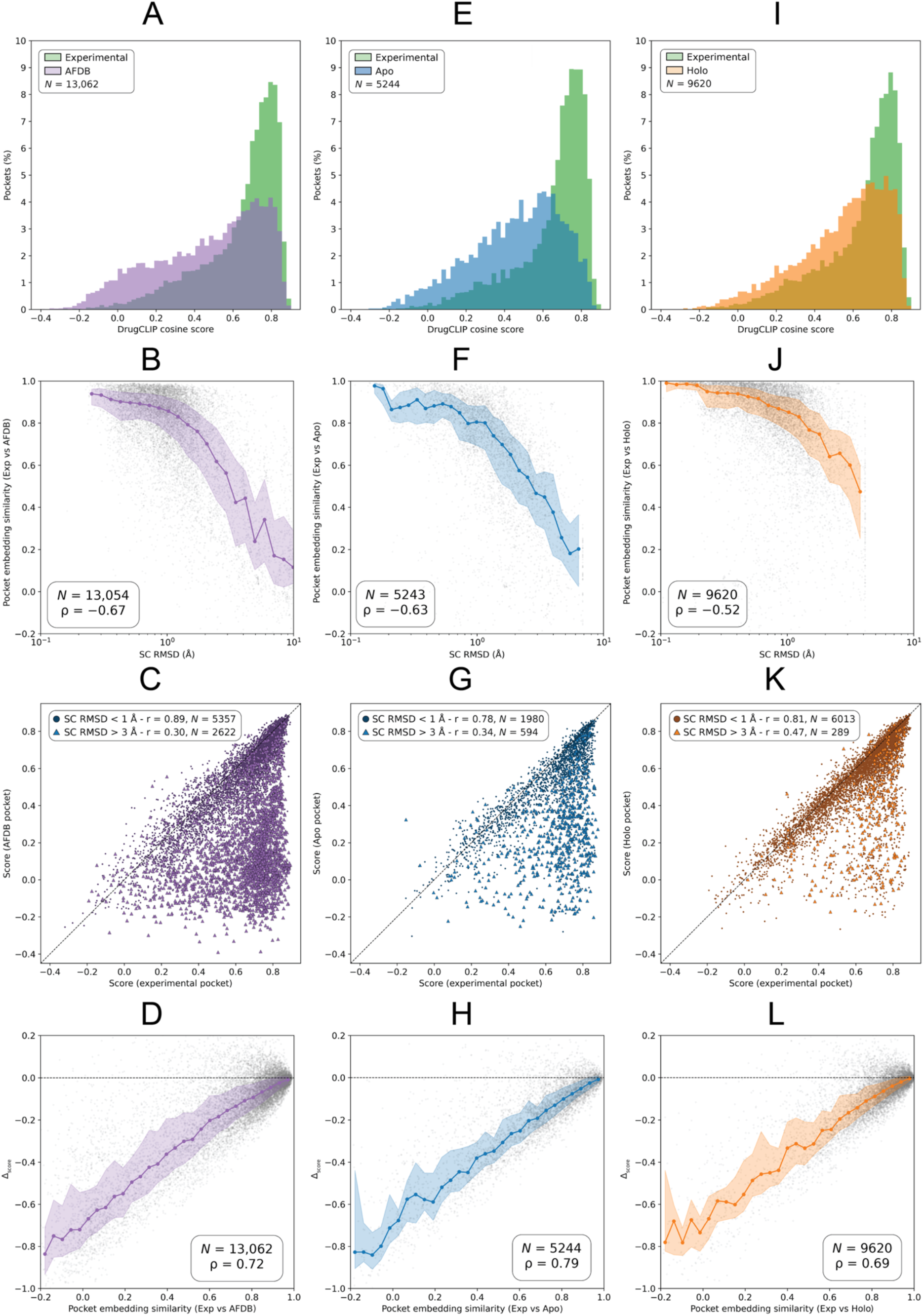
Effect of different pocket conformations on DrugCLIP scoring. Impact of alternative pocket conformations, derived from AFDB predicted models **(A-D)**, apo **(E-H)**, and holo structures **(I-L)** from AHoJ-DB have on DrugCLIP pocket-ligand screening scores, and their relationships with structural and embedding similarities. Pearson’s correlation coefficient (r) was used to assess linear relationships between screening scores, whereas Spearman’s rank correlation coefficient (ρ) was employed to evaluate monotonic associations involving structural and embedding similarities. **(A)** Distribution of pocket-ligand screening scores for 13,062 unique experimental PDB pockets (green) from the pooled data set with corresponding AFDB models, and their equivalent pockets transferred to the predicted structures (purple) – see Methods; **(B)** Relationship between sidechain (SC) RMSD and the cosine similarity between embeddings obtained from the experimental and AFDB pockets; **(C)** Scatter plot comparing DrugCLIP screening scores obtained using the experimental (X-axis) and the AFDB (Y-axis) structures. Pockets with sidechain (SC) RMSD < 1 Å are represented by dark purple circles, whereas those with SC RMSD > 3 Å are shown as lighter purple triangles. Corresponding sample sizes (*N*) and Pearson correlation coefficients (r) are indicated; **(D)** Relationship between pocket embedding cosine similarity and the change in pocket-ligand screening score (Δ_score_), calculated as AFDB minus experimental pocket score; **(E-H)** Equivalent analyses for 5244 pockets with corresponding apo conformations in AHoJ-DB; **(I-L)** Equivalent analyses for 9620 pockets with corresponding alternative holo conformations in AHoJ-DB. In panels **B**, **D**, **F**, **H**, **J** and **L**, data points displayed in grey with low opacity to reduce overplotting, and trends are represented by the median and interquartile range (IQR) within successive bins. These summaries are intended to illustrate the underlying trends and do not represent fitted regression lines or determine the reported correlation coefficients. Sample size (*N*) and Spearman’s ρ are also indicated for reference.

Figure 4 C further examines the relationship between experimental and AFDB pocket-ligand scores as a function of pocket SC RMSD, calculated following local structural alignment. Pockets with SC RMSD < 1 Å (*N* = 5357) cluster around the diagonal and exhibit a strong correlation between experimental and AFDB scores (r = 0.89). In contrast, pockets with SC RMSD > 3 Å (*N* = 2622) predominantly fall below the diagonal, indicating lower scores for AFDB pockets, and show a substantially weaker correlation (r = 0.30). These findings suggest that differences in the pocket sidechain conformations contribute to the reduced pocket-ligand similarity scores observed for AFDB structural models. Finally, Figure 4 D shows a strong positive correlation (ρ = 0.72) between experimental-AFDB pocket embedding similarity and the change in pocket-ligand score relative to the experimental structure (Δ_score_). Together, these results indicate that AFDB pockets that resemble more closely their experimental counterparts also occupy nearby regions of DrugCLIP’s embedding space and generally exhibit smaller reductions in pocket-ligand similarity scores.

Similar trends are observed for alternative apo (*N* = 5244) and holo (*N* = 9620) conformations of the canonical pockets (Figure 4 E-L). Compared with the original ligand-bound structures, 88.4% and 79.3% of pockets exhibit lower screening scores in their alternative apo and holo conformations, respectively, with corresponding median screening score differences of −0.14 and −0.05. These findings indicate that alternative holo conformations preserve better the structural features relevant to DrugCLIP’s pocket-ligand scoring than apo conformations, potentially because ligand binding favours sidechain arrangements that more closely resemble those of the canonical pocket. This is consistent with the observations of Jia *et al.* [36] and their development of GenPack to generate holo-like pocket conformations from apo structures. Furthermore, the AFDB score distribution resembles more closely that of alternative holo conformations than apo conformations, suggesting AlphaFold-predicted models are more likely to exhibit holo-like pocket geometries. This interpretation is consistent with recent findings by Saldaño *et al.* [88] and Comajuncosa-Creus *et al.* [48].

The trends observed when examining the effects of predicted protein structures and alternative apo and holo conformations on DrugCLIP scoring (Figure 4) also extend to the use of predicted binding pockets in place of experimentally determined sites, as summarised in Figure 5. Figure 5 A compares the distributions of DrugCLIP screening scores obtained using experimental PDB pockets (green), their corresponding correct P2Rank predictions (pink) – defined by DCA or DCC ≤ 4 Å – and incorrect predictions (brown), represented by the spatially closest predicted pocket for each experimental site lacking a prediction that satisfies these distance thresholds. Figure 5 B presents the equivalent analysis using P2Rank predictions obtained from the corresponding AFDB predicted models. Both panels show that the score distributions for incorrect predictions are shifted towards lower values, peaking around 0, whereas those for correct predictions peak at approximately 0.2-0.3, between the distributions for incorrect predictions and experimental pockets. These results indicate that correctly predicted pockets generally yield higher DrugCLIP screening scores than incorrect predictions, although they remain lower than those obtained using experimentally determined ligand-bound pockets.

**Figure 5.**
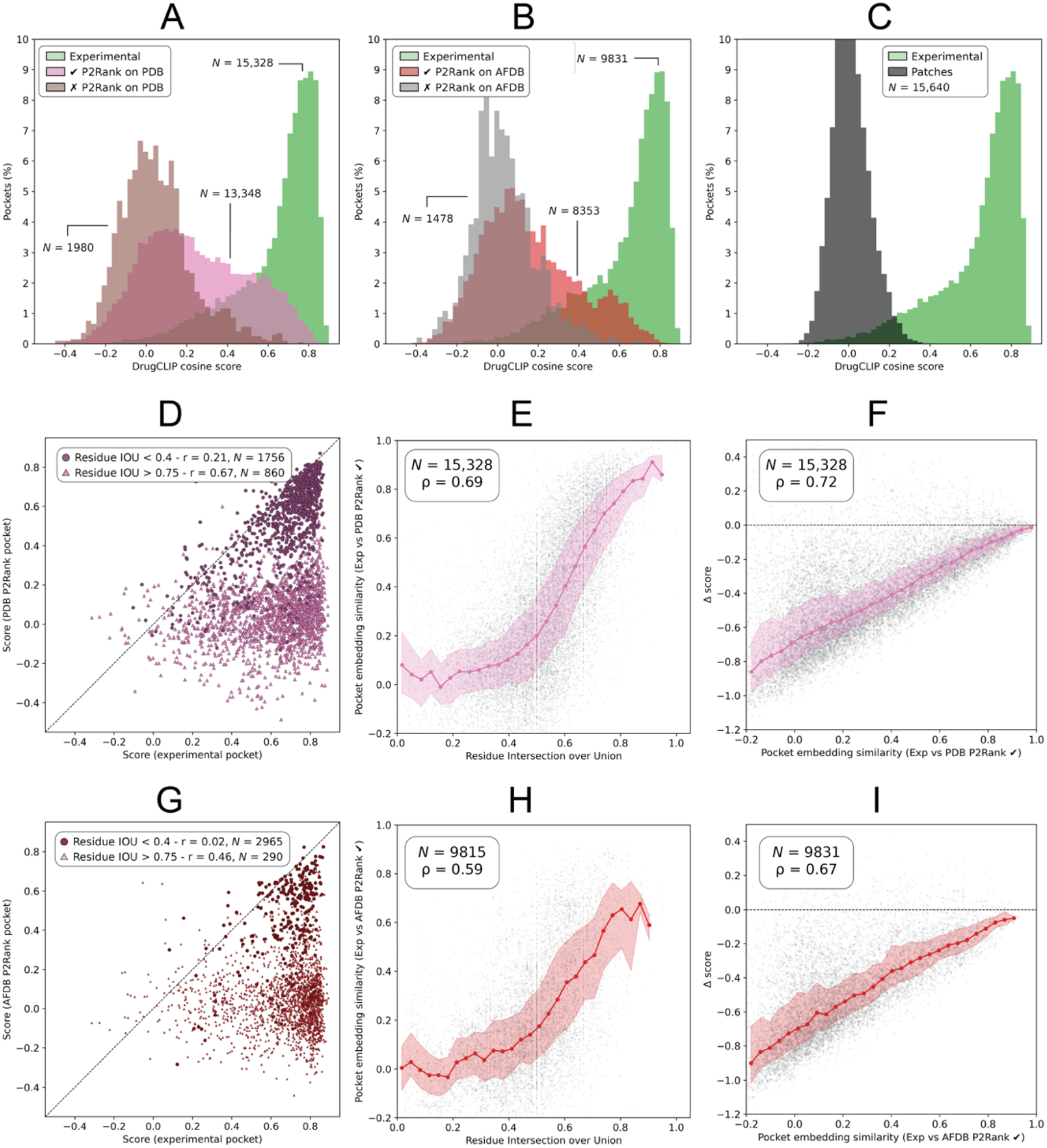
Effect of pocket prediction on DrugCLIP scoring. Impact of using P2Rank-predicted binding pockets from experimental PDB structures **(A, D-F)** or AFDB models **(B, G-I)** on DrugCLIP pocket-ligand screening scores, and their relationships with residue overlap, spatial distances and embedding similarities. Pearson’s correlation coefficient (r) was used to assess linear relationships between screening scores, whereas Spearman’s rank correlation coefficient (ρ) was employed to evaluate monotonic associations involving residue overlap, spatial distances and embedding similarities. **(A)** Distributions of pocket-ligand screening scores for experimentally determined pockets from PDB complexes (*N* = 15,328), their corresponding correct P2Rank predictions (*N* = 13,348) – satisfying either DCA or DCC ≤ 4 Å – and incorrect predictions (*N* = 1980); **(B)** Equivalent distributions for P2Rank predictions obtained from the corresponding AFDB models, comprising 9831 experimental pockets, 8353 correct predictions and 1478 incorrect predictions; **(C)** Distribution of screening scores obtained using randomly sampled protein surface patches serving as a negative control (*N* = 15,640). As expected, the resulting pocket-ligand similarity scores are centred around zero; **(D)** Scatter plot comparing DrugCLIP screening scores obtained using experimental (X-axis) and predicted (Y-axis) pockets. Predictions with a residue intersection over union (IOU) > 0.75 are represented by dark purple circles, whereas those with IOU < 0.4 are shown as lighter purple triangles. Corresponding sample sizes (*N*) and Pearson correlation coefficient (r) are indicated; **(E)** Relationship between residue overlap (IOU) and pocket embedding cosine similarity; **(F)** Relationship between pocket embedding cosine similarity and the change in pocket-ligand screening score (Δ_score_), calculated as predicted minus experimental pocket score; **(G-I)** Equivalent analyses to **D-F** using P2Rank predictions obtained from AFDB models instead of experimental PDB structures.

Figure 5 C shows the distribution of DrugCLIP screening scores obtained by evaluating 300,587 random protein surface patches against the corresponding bound ligands across 15,640 pockets (see Methods). These random patches serve as a negative control for assessing the similarity between DrugCLIP’s learned pocket and ligand representations in the absence of a known binding site. Compared with the distribution of incorrectly predicted pockets, the random surface patch distribution exhibits a taller peak around 0 (12.5%) and a narrower spread, with 95% of scores falling between –0.17 and 0.19. Its tails are also considerably less pronounced, with a maximum observed score of 0.45.

Figure 5 D-I and Figure S2 explore the relationship between DrugCLIP embedding similarity for experimental and predicted pockets and four criteria used to classify pocket predictions as correct (true positives) or incorrect (false positives) relative to the experimentally observed binding sites. Two criteria are distance-based: distance between centres (DCC) and distance between the predicted pocket centre and the nearest atom of the observed ligand (DCA). The remaining two assess residue overlap: residue recall and intersection over union (IOU). Residue recall measures the proportion of experimentally observed pocket residues included in the predicted pocket. A large predicted pocket could achieve perfect recall simply by encompassing a smaller experimental pocket, regardless of the additional residues included. To account for this limitation, residue IOU was also considered, as it measures the overlap relative to the combined set of residues in both pockets.

Figure 5 E, H and Figure S2 A-C, G-I show that embedding cosine similarity between experimental and predicted pockets correlates more strongly with residue IOU (ρ_P2Rank PDB_ = 0.69; ρ_P2Rank AFDB_ = 0.59), than with DCC (ρ_P2Rank PDB_ = –0.62; ρ_P2Rank AFDB_ = –0.49), DCA (ρ_P2Rank PDB_ = –0.33; ρ_P2Rank AFDB_ = –0.27) or residue recall (ρ_P2Rank PDB_ = 0.44; ρ_P2Rank AFDB_ = 0.09). This suggests that DrugCLIP’s pocket embeddings are more sensitive to differences in residue membership and overlap than to the spatial proximity of predicted and experimental pocket centres. Figure 5 D and G show substantially stronger correlations between experimental and predicted pocket-ligand screening scores for predictions with high (> 0.75) residue IOU (r_P2Rank PDB_ = 0.67; r_P2Rank AFDB_ = 0.46) than for those with low (< 0.4) IOU: (r_P2Rank PDB_ = 0.21; r_P2Rank AFDB_ = 0.02). Differences in score correlations are less pronounced when predictions are grouped according to residue recall, DCA or DCC (Figure S2 D-F, J-L) – although DCC shows the strongest association with embedding similarity among these three criteria.

Finally, consistent with the trends observed for alternative pocket conformations (Figure 4 D, H, L), predicted pockets with greater embedding cosine similarity to their experimental counterparts generally exhibit smaller reductions in pocket-ligand screening scores (Δ_score_) relative to the experimentally observed complexes (Figure 5 F, I).

### Performance robustness and degradation analysis

Building on the preceding analyses, Figure 6 summarises how predicted protein structures, alternative apo and holo conformations, and pocket prediction affect DrugCLIP’s screening scores and how these changes translate into bound ligand recovery performance. Specifically, we investigate how performance deteriorates under increasingly realistic screening conditions that progressively depart from the ideal scenario in which both the experimentally determined binding pocket structure and the crystallographic ligand pose are known.

**Figure 6.**
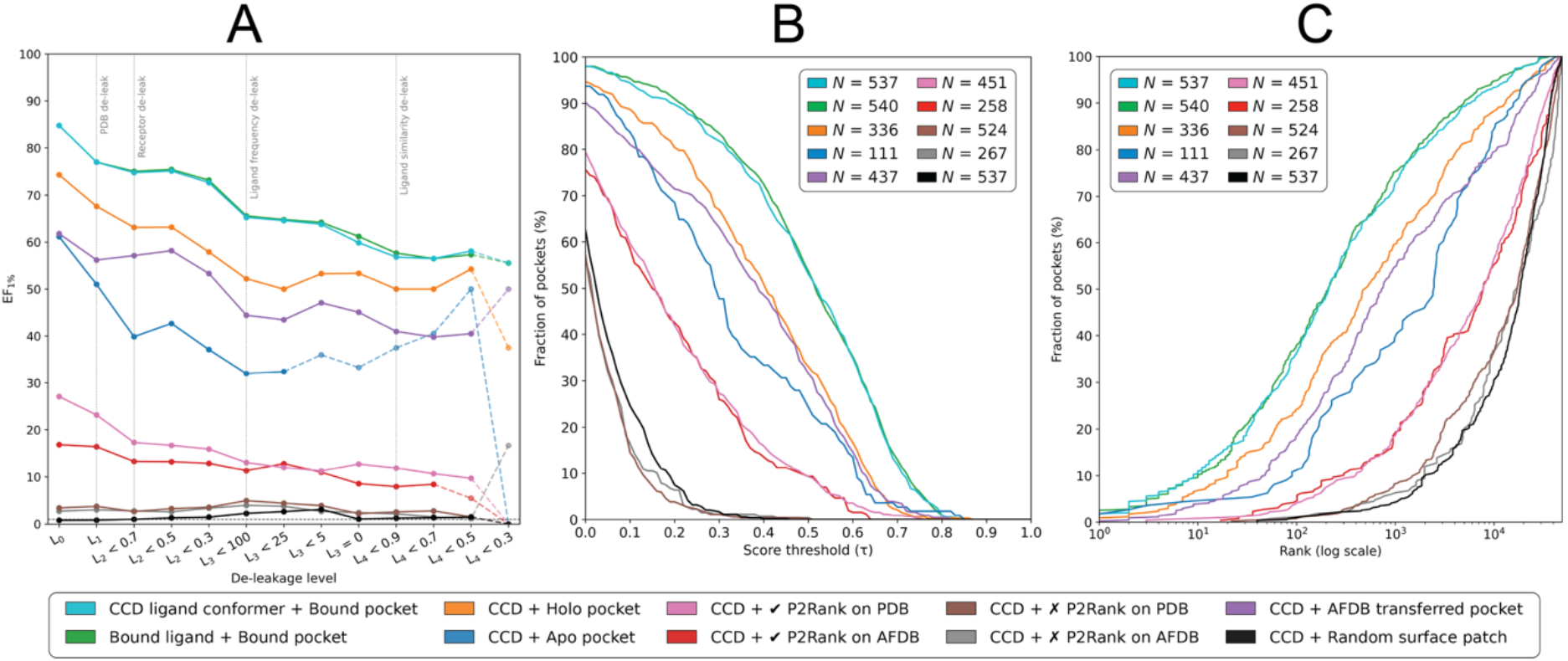
DrugCLIP performance degradation analysis. Summary of DrugCLIP’s performance degradation on the bound ligand recovery benchmark across increasingly challenging subsets of pocket-ligand pairs, progressively less similar to the BioLiP subset used to fine-tune DrugCLIP. Performance was also evaluated under increasingly realistic virtual screening conditions, including alternative experimentally determined apo and holo conformations, AFDB predicted models, and P2Rank-predicted pockets from both experimental PDB structures and AFDB models. Three negative controls are included for reference: incorrectly predicted pockets from PDB structures and AFDB models, and randomly sampled protein surface patches. See Methods for further details. **(A)** Enrichment factor at 1% (EF_1%_) across successive de-leakage levels for CCD ligand conformers screened against experimental bound pockets (cyan), crystallographic ligand conformers against experimental bound pockets (green), CCD conformers against holo pockets (orange), apo pockets (blue), AFDB pockets (purple), correctly predicted P2Rank pockets from PDB structures (pink) and AFDB models (red), incorrectly predicted P2Rank pockets on PDB structures (brown) and AFDB models (grey), and random surface patches (black). Vertical light grey dotted lines mark the boundaries between successive de-leakage criteria: same PDB overlap, protein sequence identity, ligand occurrence frequency, and ligand chemical similarity (ECFP4 TS) relative to the fine-tuning set. Coloured dashed lines indicate subsets containing fewer than 100 evaluated pockets. The number of pockets evaluated under each condition and de-leakage level is reported Table S5; **(B)** Score survival curves showing the fraction of pockets which cosine similarity score with the experimentally observed bound ligand exceeds a given threshold. A cosine similarity threshold of 0.5 was selected as an illustrative reference point, corresponding the upper quartile of the theoretical similarity range (−1 to 1). Whereas a score of 0 indicates orthogonal embeddings, a score of 0.5 corresponds to an angle of 60° between vectors, indicating positive directional alignment. Although this threshold does not represent a calibrated measure of binding compatibility, it provides a consistent reference for comparing pocket-ligand embedding similarities across screening conditions. The X axis is truncated at 0 for clarity; **(C)** Cumulative proportion of pockets as a function of the rank of their experimentally observed bound ligand among the 48,999 compounds in the CCD screening library. Panels B and C use the de-leakage level L_3_ < 25, the most stringent level at which all screening conditions retain at least 100 pockets.

As in Figure 3 and Table 2, Figure 6 A illustrates how DrugCLIP’s enrichment factor at 1% (EF_1%_) changes as the initial pooled set of 15,533 pockets is progressively restricted through a de-leakage analysis. This sequential filtering considers protein sequence identity, ligand frequency and ligand chemical similarity relative to DrugCLIP’s BioLiP fine-tuning set, yielding increasingly challenging pocket-ligand pairs. All screening conditions, except the three negative controls, exhibit a similar trend, with EF_1%_ decreasing as the evaluated pockets become less similar to those encountered during fine-tuning. The negative controls comprise CCD ligand conformers screened against incorrectly predicted P2Rank pockets from PDB structures or AFDB models, and randomly sampled protein surface patches. Across the seven remaining conditions, EF_1%_ decreases from 84.8, 84.8, 74.3, 61.2, 61.8, 27.1 and 16.4 at de-leakage level 0 (L_0_ – full set) to 64.6, 64.8, 50.0, 32.4, 43.5, 12.0 and 12.8 at L_3_ < 25, respectively. These vales correspond, in order, to CCD ligand conformers screened against experimental bound pockets, bound ligand conformers against experimental bound pockets, and CCD conformers against alternative holo, apo and AFDB pockets, followed by correctly predicted P2Rank pockets from PDB structures and AFDB models. At L_3_ < 25, complexes sharing structures with the fine-tuning set, proteins with > 30% sequence identity and ligands observed ≥ 25 times during fine-tuning have been excluded. We focus on this level because it represents the most stringent de-leakage threshold at which all screening conditions retain at least 100 evaluable pockets.

Figure 6 B further examines the impact of these conditions on the distributions of pocket-ligand embedding cosine similarity scores. A threshold of 0.5 was selected as an illustrative reference point to facilitate comparisons between distributions, rather than as a criterion for successful ligand retrieval, which depends on the relative ranking of candidate compounds. Although a reduction in absolute pocket-ligand similarity does not necessarily imply poorer retrieval, Figure S4 reveals strong negative correlations between changes in cosine similarity and retrieval rank across screening conditions (ρ = −0.79 to −0.91), indicating that lower scores are generally associated with poorer ligand retrieval. The nearly overlapping distributions obtained by using crystallographic and CCD ligand conformers further demonstrate DrugCLIP’s robustness to ligand conformational variation, with 53.7 and 53.1% of pockets achieving scores ≥ 0.5 (P_s ≥ 0.5_), respectively. Furthermore, the AFDB score distribution resembles more closely that of alternative holo conformations than apo conformations, consistent with the trends observed in Figure 4. The percentages of pockets achieving scores ≥ 0.5 are 24.3, 33.0 and 31.6% for apo, holo and AFDB pockets, respectively.

Finally, Figure 6 C illustrates how these differences translate into bound ligand retrieval performance through the cumulative distributions of ligand ranks within the 48,999-compound CCD library. The median rank increases from 212 for CCD ligand conformers screened against experimental bound pockets to 494, 733 and 2470 for alternative holo, AFDB and apo conformations, respectively. Performance deteriorates further when predicted pockets are used, with median ranks of 7753 and 7801 for P2Rank predictions obtained from PDB structures and AFDB models, respectively. These results highlight the importance of accurately identifying the ligand-binding residues and preserving their local structural arrangements, particularly sidechain orientations, for effective DrugCLIP screening, consistent with the findings of Jia *et al.* [36]. A detailed breakdown of these results is presented in Table 3 and Tables S5 and S6.

**Table 3.**
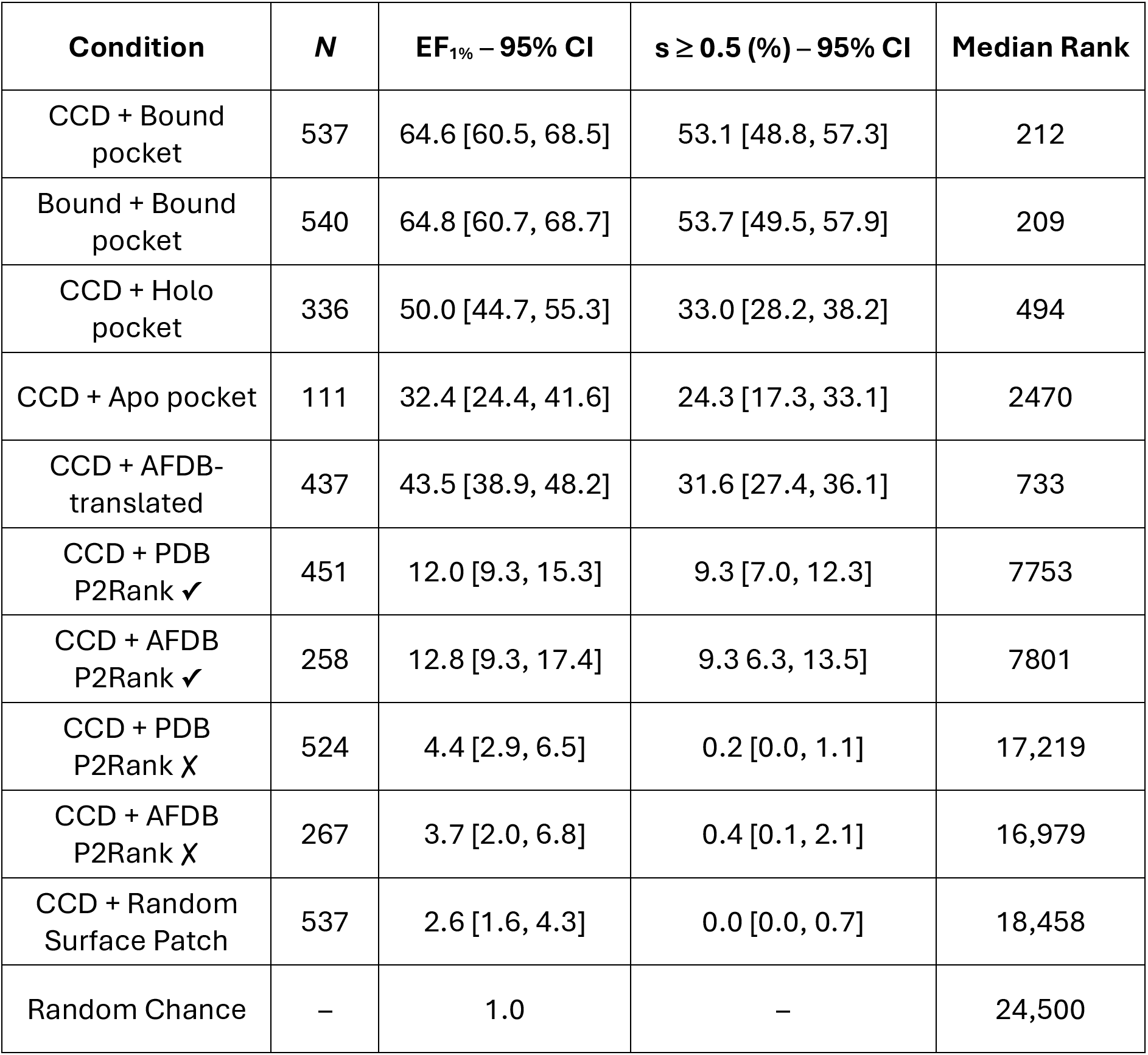
Summary of DrugCLIP performance degradation at de-leakage level L_3_ < 25. Summary of DrugCLIP’s performance under nine screening conditions designed to reflect more closely realistic structure-based virtual screening scenarios, alongside a random retrieval baseline for reference. For each condition, the number of evaluated pockets (*N*), enrichment factor at 1% (EF_1%_), percentage of pockets achieving a protein-ligand screening score ≥ 0.5 (%) and median rank of the experimentally observed bound ligand are reported. Values in brackets indicate 95% confidence intervals for EF_1%_ and s ≥ 0.5 (%).

**Table 4.**
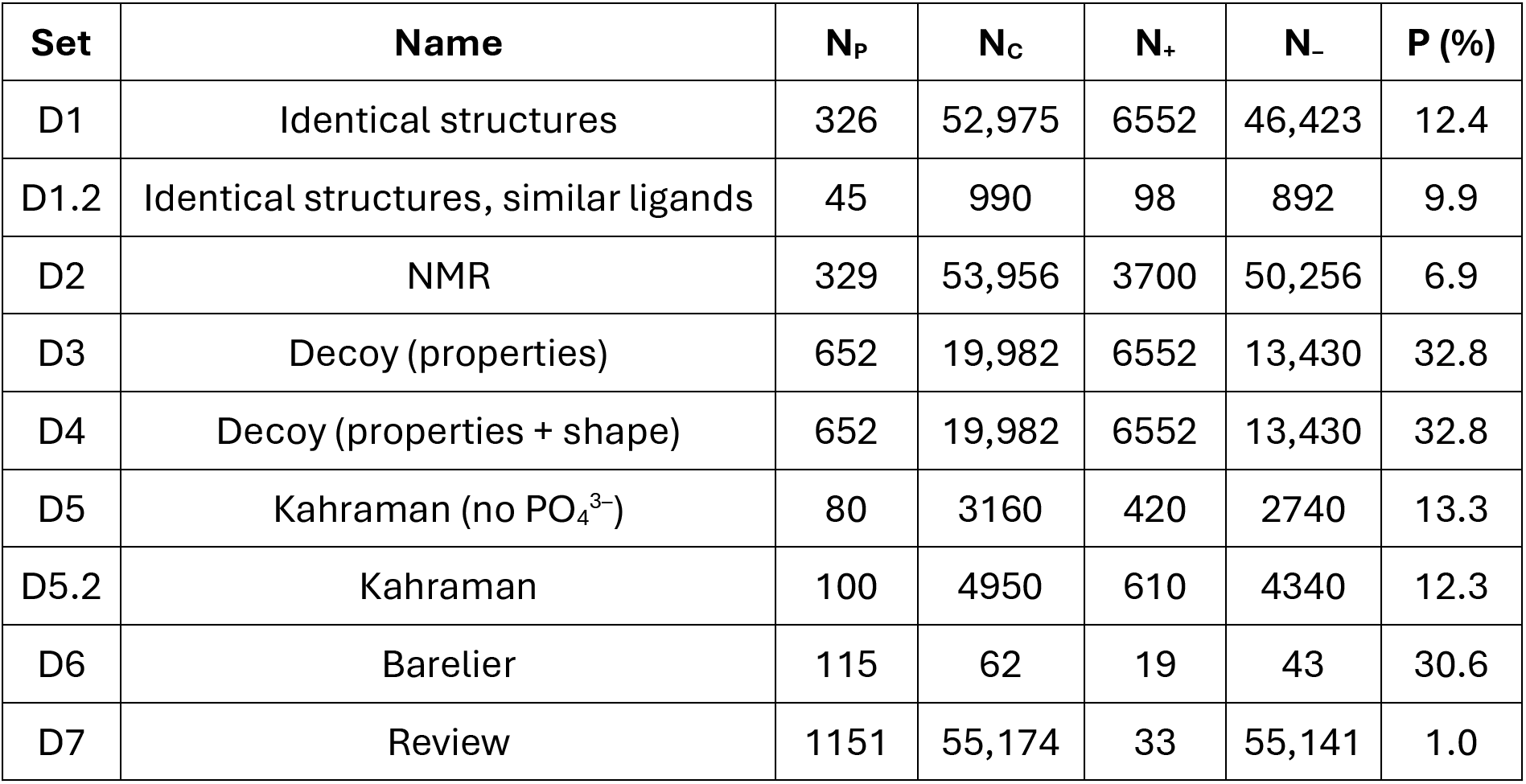
ProSPECCTs data sets summary. Summary of the 9 ProSPECCTs sets used in our pocket matching benchmark. D6.2 is excluded since cofactor atoms are not valid for DrugCLIP’s pocket encoder. N_P_ is the total number of pockets in the set. N_C_ is the number of comparisons or pocket pairs in the set, of which N_+_ are active, similar or positive pairs, representing a percentage P (%) of N_P_, and N_–_ are inactive, dissimilar or negative pairs. N_C_, N_P_, and N_–_ differ from the figures in the original article by Ehrt *et al.* [45] because only unique non-identical pairs were considered in this benchmark.

As the availability of alternative pocket representations varies across proteins, the number of pockets evaluated differs between experimental conditions. To determine whether these differences affect our conclusions, we repeated the de-leakage analysis using only pockets shared between the conditions being compared (Figure S3). Both EF_1%_ values and overall trends remained largely consistent with those obtained from the original analyses, suggesting that differences in pocket coverage do not materially influence our conclusions. Nevertheless, comparisons at the strictest de-leakage levels should be interpreted cautiously given the small number of remaining pockets.

## Discussion

This study presents a comprehensive series of analyses and benchmarks that complement the original work of Gao *et al.* [35] and Jia *et al.* [36], providing a clearer understanding of what DrugCLIP’s encoder modules learn. Our findings shed light on the information captured by the abstract representations of protein pockets and ligands, revealing aspects of the structure and organisation of the latent space learned during model training. Furthermore, by evaluating DrugCLIP across multiple tasks and conducting a systemic de-leakage analysis, we investigate the extent to which its performance depends on memorisation of the fine-tuning set data versus generalisation beyond them, while identifying its strengths and limitations under different screening conditions.

In this study, we systematically evaluated DrugCLIP’s pocket matching capabilities by comparing its performance against established methods for pocket structural comparison and similarity search tools, as well as more recently developed pocket descriptors such as EPoCS [46] and PocketVec [48], across the ProSPECCTs data sets. DrugCLIP achieved the highest average classification performance (AUC = 0.87), establishing a new state of the art in pocket similarity search. Remarkably, DrugCLIP achieved this performance despite not being explicitly trained for pocket matching, but rather to distinguish binding from non-binding molecules in a virtual screening context. Nevertheless, its training objective appears to have encouraged the development of pocket representations that capture information relevant not only to identifying compatible ligands but also to recognising structurally similar binding pockets. Furthermore, DrugCLIP offers a speed improvement of over 10,000-fold compared with some existing pocket descriptors, making it a particularly promising approach for large-scale pocket characterisation and similarity searches across the protein universe.

A comprehensive analysis of DrugCLIP’s learned representations was conducted using data sets of protein-ligand complexes commonly employed to train, validate and evaluate ligand binding site prediction methods. To investigate the information encoded within these abstract representations, we compared embeddings generated from different conformers of the same molecule and examined the relationships between embedding similarities and conventional measures of chemical and structural similarity, including the Tanimoto coefficient calculated from extended connectivity fingerprints (ECFP4) and root mean square deviation (RMSD). Our findings demonstrate DrugCLIP’s robustness to ligand conformational variation and reveal a strong correspondence between pocket embedding similarity and structural similarity, further supporting the results of our pocket matching benchmark. In contrast, although ligand embedding similarity is positively correlated with chemical fingerprint similarity, this relationship is considerably weaker than that observed between pocket embeddings and their structural counterparts. One possible explanation is that DrugCLIP’s contrastive learning objective encourages its latent representations to capture aspects of ligand compatibility with binding environments beyond those reflected by conventional chemical descriptors. Consequently, chemically dissimilar molecules may be positioned close together in the ligand embedding space if they share features relevant to binding similar pockets, despite exhibiting limited similarity according to their molecular fingerprints. This interpretation is supported by the stronger correlation between ligand embedding similarity and conventional chemical similarity observed for ligands associated with more similar binding pockets.

DrugCLIP’s ability to recover the experimentally observed bound ligand was evaluated by screening a library of 48,999 compounds from the CCD, including the crystallographic ligand, against a set of unique pockets pooled from the COACH420, HOLO4K, scPDB and PDBBind data sets. For each pocket, the rank of the bound ligand within the library was recorded, and performance was assessed using enrichment factor at 1% (EF_1%_), among other metrics. Performance was then examined across progressively more restrictive subsets of pocket-ligand pairs, obtained through a sequential de-leakage analysis designed to reduce similarity to DrugCLIP’s fine-tuning set. Our results provide evidence that DrugCLIP can generalise beyond its fine-tuning data to previously unseen proteins and ligands, ranking the crystallographic ligand within the top 1% of molecules for 56.5% of pockets associated with proteins sharing ≤ 30% sequence identity and ligands exhibiting ≤ 0.7 chemical similarity to those in the fine-tuning set. Nevertheless, although our sequential filtering strategy accounts for multiple sources of training-test similarity, proteins with low sequence identity may still share structurally similar ligand binding pockets. Therefore, we cannot entirely exclude the possibility that similarities between binding environments represented in the fine-tuning and evaluation sets contribute to the observed performance.

Furthermore, we evaluated DrugCLIP’s robustness under several conditions that resemble more closely realistic virtual screening scenarios, in which the protein-ligand complex is unknown and only alternative experimentally determined apo or holo structures, predicted protein structure models or computationally identified binding pockets are available. Our results reveal a smaller performance decrease when using AFDB pocket conformations than alternative apo conformations, although performance remains below that achieved with experimental holo structures. This suggests that AlphaFold-predicted models tend to exhibit pocket geometries more closely resembling holo than apo conformations, in agreement with previous studies [48, 88]. In these comparisons, pocket residue membership is preserved, with only atomic coordinates varying between structures. Consequently, the observed performance degradation primarily reflects conformational differences, particularly changes in sidechain orientation, rather than inaccuracies in pocket residue selection. Importantly, these analyses rely on pocket residues initially defined from experimentally determined holo complexes and subsequently transferred onto alternative structures. Therefore, the performance observed with AFDB models should be regarded as an optimistic estimate for scenarios in which the binding site location and residue composition are already known. Conversely, comparing experimentally defined and P2Rank-predicted pockets on the same protein structure allows us to isolate the effects of pocket residue selection independently of protein conformational variation. The resulting performance differences are strongly associated with the intersection over union between predicted and experimentally observed pocket residues. In fully *de novo* virtual screening applications, where binding pockets must also be identified computationally, further performance degradation is observed, highlighting the importance of accurate pocket identification alongside the preservation of appropriate pocket conformation.

These findings highlight the importance of accurately identifying ligand-binding residues and preserving their sidechain orientations to maximise DrugCLIP’s performance. The importance of sidechain placement had previously been recognised by Jia *et al.* [36], who developed GenPack, a generative model conditioned on pocket backbone atoms that refines sidechain conformations to improve DrugCLIP’s screening performance. Although this approach can help reduce conformational differences between alternative apo and holo structures of the same binding pocket, it does not directly address discrepancies in residue composition between experimentally observed and predicted pockets. In light of the findings reported by Utgés and Barton [74], we anticipate that binding site prediction methods achieving greater residue and volume overlap with experimentally observed pockets, such as PUResNet [89], GrASP [73] and DeepPocket [90], could further improve the performance of DrugCLIP and similar approaches compared with predictions obtained using tools such as fpocket [91] or P2Rank [72].

Although our study focuses exclusively on DrugCLIP, the analytical approach presented here could be extended to other methods such as DrugHash [29], BindCLIP [44], or ConGLUDe [41], which similarly encode protein pockets and ligands as vectors representations for virtual screening and related tasks. In particular, these analyses could help investigate potential limitations of contrastive learning approaches, including shortcut learning, insufficient sensitivity to fine-grained atomic interactions, and variable performance across protein families. Comparing embedding similarities with established measures of structural and chemical similarity, alongside appropriate negative controls, could help determine whether the learned representations capture meaningful molecular features or are influenced by superficial correlations. Likewise, assessing the effects of conformational changes and pocket definitions, together with stratifying screening performance by protein family, could reveal limitations in capturing specific molecular interactions or generalising across different binding environments. Overall, our findings provide insight into the representations learned by DrugCLIP, highlighting both their strengths and limitations, while offering a conceptual framework for systematically evaluating and guiding the development of future contrastive learning methods for structure-based virtual screening.

### Conclusions

The conclusions resulting from this work are as follows:

- DrugCLIP achieves state-of-the-art performance in pocket similarity search, outperforming dedicated descriptors despite not being explicitly trained for this task, while offering a speed advantage of over 10,000-fold compared with existing methods.
- DrugCLIP’s latent representations capture meaningful structural and chemical information: pocket embeddings strongly reflect structural similarity, while ligand embeddings capture aspects of molecular similarity beyond conventional chemical descriptors, potentially reflecting compatibility with similar binding environments.
- DrugCLIP demonstrates generalisation beyond its fine-tuning set, ranking the crystallographic bound ligand in the top 1% of 48,999 compounds for 56.5% of pockets associated with previously unseen proteins and chemically dissimilar ligands (*N* = 223).
- Performance deteriorates under increasingly realistic screening conditions, with sidechain conformational differences contributing to degradation when using alternative holo, apo, and AFDB structures, and inaccurate residue selection affecting performance with predicted pockets.
- Accurate pocket identification and sidechain placement are critical to DrugCLIP’s performance. Pocket predictors achieving higher residue and volume overlap with experimentally observed pockets, such as PUResNet and GrASP, could further improve screening performance.
- Our analytical framework provides a foundation for investigating the learned representations and generalisation capabilities of other contrastive learning pocket-ligand models, including BindCLIP and ConGLUDe.

## Methods

### Pocket matching benchmark

The ability of DrugCLIP’s pocket encoder to perform pocket matching was evaluated using the ProSPECCTs (Protein Site Pairs for the Evaluation of Cavity Comparison Tools) benchmark suite [45]. ProSPECCTs comprises ten data sets of protein pocket or cavity pairs, each designed to assess pocket similarity under different contexts and scenarios. In these sets, pocket pairs are labelled as *actives* (“similar”) or *inactives* (“dissimilar”) according to different criteria with the aim of capturing binding site similarity at different levels. In this way, each of these sets covers different definitions of binding pocket similarity.

Dataset 1 (D1) consists of 326 protein-ligand complexes distributed across 12 groups of structures with identical protein sequences but different bound ligands. This set evaluates the sensitivity of pocket comparison methods to differences in binding site definition arising from the binding of different ligands to the same pocket. D1.2 is a subset of D1 restricted to identical or similar ligands, comprising 45 complexes across the same 12 proteins. In both data sets, pocket pairs belonging to the same protein are considered similar, whereas all other pairs are considered dissimilar. D2 assesses the influence of binding site flexibility by comparing different pocket conformations sampled from 329 NMR models across 17 protein structures. Pocket pairs belonging to the same NMR ensemble are classified as similar, and all others as dissimilar. D3 and D4 are decoy data sets, generated by introducing one to five artificial mutations into the original D1 pockets, altering their physicochemical properties (D3) or both their physicochemical properties *and* shape (D4). For consistency with previous studies by Ehrt *et al.* [45] and Comajuncosa-Creus *et al*. [48], only the subset including five mutations was considered, which contains 326 of the 1630 mutated structures. In these two sets, original (unmutated) pocket pairs belonging to the same protein are labelled as similar, whereas all comparisons between wild-type and mutated structures are classified as dissimilar. D5.2 contains 100 evolutionarily diverse proteins, belonging to different CATH [92, 93] homologous superfamilies, including 20 phosphate binding sites, collated by Kahraman *et al*. [49]. This set assesses the ability of pocket comparison methods to recognise evolutionarily distinct sites that bind the same ligand as similar. D5 is a subset of D5.2 that excludes the 20 phosphate sites, resulting in 80 pockets. In both sets, pocket pairs binding the same ligand are classified as similar, whereas all others as dissimilar. D6, compiled by Barelier *et al.* [50], includes 116 protein-ligand complexes involving 59 different ligands, yielding 62 pocket pairs from evolutionary diverse proteins according to PFAM [94, 95], SCOP [96, 97] and CATH [98, 99] classifications. These pairs are labelled as similar (class A) when the same ligand groups interact with similar protein environments, or dissimilar when the same ligand groups interact with different protein environments (class B) or different ligand groups interact with the protein (class C). D6.2 contains the same set of structures as D6 but retains additional bound cofactors that were excluded from the original data set, affecting 22 of the 116 complexes. D6.2 was excluded from the present benchmark because these additional ligand atoms would not be correctly processed by DrugCLIP’s pocket encoder. Finally, D7 contains 1151 structures and 115 known similar binding site pairs reported in the literature [51], encompassing both proteins from the same family, as well as those belonging to unrelated families. All remaining pocket pairs are dissimilar. Evaluation was performed on unordered pocket pairs, excluding self-comparisons, i.e., A *vs* B, but not B *vs* A, A *vs* A or B *vs* B. Dataset size and label frequencies are reported on Table 4.

Out of the 9 sets evaluated, D1.2 is expected to be the least challenging, as both the pocket sequences and bound ligands are identical or highly similar. D1 would follow in order of increasing difficulty. D2 should be relatively straightforward, as positive pairs correspond to different conformations of the same binding pocket within the same protein, unless substantial structural differences exist between NMR models. Performance on D5 should be better than D5.2, as the former excludes phosphate binding sites, which are ubiquitous and structurally diverse due to the small size of the phosphate group. D3, D4, D6 and D7 are expected to be the most challenging, as their similarity labels are based on artificially induced mutations, differences in ligand binding modes or a combination of similarity criteria. However, D3 and D4 warrant particular caution, as their mutations were introduced through sidechain replacement without any reported subsequent structural relaxation or energy minimisation, potentially yielding physically unrealistic binding site conformations. Furthermore, these mutations were not experimentally validated to alter ligand recognition, raising questions about the biological relevance of the resulting similarity labels.

In their original study, Jia *et al.* [36] evaluated the ability of DrugCLIP’s pocket encoder across several downstream tasks, including pocket matching, using ProSPECCTs D5, originally collated by Kahraman *et al.* [49] and excluding phosphate binding sites. Here, we extend this evaluation to the remaining ProSPECCTs data sets and compare DrugCLIP’s performance against other recently developed methods for ligand binding site representation and characterisation, including EPoCS [46] and PocketVec [48], the structure-based virtual screening method NRGRank [47], as well as additional DrugCLIP variants and relevant baselines. Discriminatory performance, or classification accuracy, is assessed using the area under the receiver operating characteristic curve (AUROC), consistent with previous studies [36, 45, 48].

EPoCS (ESM-driven Pocket Cross-Similarity) is a metric to quantify similarity between protein binding sites [46]. EPoCS employs radially truncated Voronoi tessellation [100, 101] around a bound ligand to identify interacting protein atoms and their corresponding residues, thereby defining the binding pocket. Each pocket residue is represented by an embedding generated using the ESM-2 [10] protein language model, and these residue-level embeddings are pooled into a single vector representing the entire pocket. Pocket dissimilarity is then quantified as the Euclidean distance between the corresponding pocket vectors.

PocketVec generates a pocket representation that captures the chemical environment and ligand binding landscape of a binding site by docking a library of 128 selected lead-like molecules (LLMs) using rDock [50]. The resulting descriptor is a vector containing the ranks of these molecules based on their docking scores. As with EPoCS, pocket dissimilarity is quantified using the Euclidean distance between the corresponding descriptor vectors [48].

NRGRank is a coarse-grained, structurally informed method designed for ultra-large-scale virtual screening [47]. NRGRank evaluates pairwise protein-ligand atom type interactions using a pseudo-energy function that accounts for ligand and sidechain flexibility, as well as limited backbone movement. For each protein-ligand complex, multiple ligand conformations are sampled within the binding pocket grid, and the conformation with the most favourable pseudo-energy is selected to determine the NRGRank score. Here, we adapted the PocketVec approach by screening the same library of 128 lead-like molecules against the ProSPECCTs pockets using NRGRank, thereby, repurposing this virtual screening method to generate pocket descriptors.

For DrugCLIP, pocket similarity was quantified using the cosine similarity between pocket embeddings, following Jia *et al.* [36]. DrugCLIP was employed using its publicly available pretrained model (https://github.com/THU-ATOM/Drug-The-Whole-Genome) and default inference configuration, without additional post-processing of the resulting 128-dimensional embeddings. Three additional variants were also surveyed. DrugCLIP_12 Å_ defines pockets by including all residues with at least one atom within 12 Å of the bound ligand, rather than the default 6 Å threshold. DrugCLIP_Sim Vec_ adapts the PocketVec approach by leveraging DrugCLIP’s virtual screening capabilities to generate pocket descriptors. Specifically, the same library of 128 lead-like molecules was encoded using DrugCLIP’s molecule encoder, and cosine similarities between each pocket and the 128 ligand embeddings were calculated to generate a vector of screening scores. These similarity scores were subsequently converted into ranks, yielding the third variant: DrugCLIP_Rank Vec_.

To provide a broader comparison across different methodological approaches, we also considered 21 additional pocket matching methods, comprising the sixteen methods [53–66] previously benchmarked by Ehrt *et al.* [45] and five more recently developed approaches: DeeplyTough [67], Site2Vec [68], BindSiteS-CNN [69], SiteMine [70] and SiteContext [71]. These methods encompass a range of strategies for representing and comparing binding sites, including geometric alignment, pharmacophore and fingerprint descriptors, and machine learning-based representations. Their published AUROC values on the ProSPECCTs data sets, obtained from Ehrt *et al.* [44] and the respective original studies [67–71] were used for comparison with the methods evaluated here.

Finally, several naïve baselines were included for comparison. Two binary baselines assign similarity labels based on whether pocket pairs belong to the same protein, as determined by their UniProt accession [102, 103] – or 100% sequence identity (SI) for D3 and D4 – or bind the same ligand, as determined by their Chemical Component Dictionary (CCD) identifier [86]. Two additional baselines use pocket sequence identity, calculated following global sequence alignment, and ligand chemical similarity, quantified using the Tanimoto coefficient (TC) [104–106] between extended connectivity fingerprints (ECFP4) – 2048 bits, radius 2 – [107, 108] generated with RDKit v2026.03.02 [109]. MMseqs [110–112] was used for pairwise sequence alignment and sequence identity calculations. Importantly, these baselines should not be regarded as minimum performance thresholds, as some can achieve perfect discrimination power by directly reflecting the criteria used to assign similarity labels. For example, sequence identity can perfectly distinguish similar from dissimilar pairs in D1, D1.2 and D2, where positive pairs correspond to pockets from the same protein, as well as in D3 and D4, where artificially introduced mutations distinguish negative pairs from their wild-type counterparts. Likewise, ligand identity or chemical similarity can provide strong discriminatory power for D5 and D5.2, where pocket similarity is defined according to whether evolutionarily distinct proteins bind the same ligand.

### Protein-ligand complexes data sets

Four data sets commonly used for training, validation and testing of ligand binding site prediction methods were employed to evaluate DrugCLIP’s ability to recover experimentally bound ligands and to investigate its learned pocket and molecule representations. These correspond to the subsets of COACH420 [113], HOLO4K [77, 114], scPDB [115, 116] and PDBBind (2020) [117, 118] used by Sestak *et al.* [75] in their recent work (https://zenodo.org/records/17365855) [119]. COACH420 contains 420 single-chain protein structures bound to a variety of drug-like molecules and natural ligands. HOLO4K is a larger set of approximately 4000 structures including both single-and multi-chain complexes (https://github.com/rdk/p2rank-datasets). Consistent with previous studies [75, 90, 120], the “Mlig” subsets of both data sets were employed, retaining biologically relevant ligands, as defined by the binding Mother of all Databases (MOAD) [121, 122]. These subsets include 265 chains and 342 binding sites for COACH420 and 4639 chains and 6744 sites for HOLO4K. The number of sites can exceed the number of protein chains because multiple ligands may bind at distinct locations within the same protein. scPDB is a comprehensive database of pharmacologically relevant protein-ligand complexes. The subset of 5019 structures utilised here was obtained by clustering the 16,034 entries in the 2017 release by sequence, following Sestak *et al.* [75] and Kandel *et al.* [123]. PDBBind provides an extensive collection of experimentally measured binding affinity data for biomolecular complexes (https://www.pdbbind-plus.org.cn/download). Here, we used the refined set from its 2020 release, which includes 5316 high-quality curated entries. For scPDB and PDBBind, only one ligand was retained per structure, resulting in an equal number of protein structures and binding pockets.

### DrugCLIP representations’ analysis

Multiple experiments were conducted to investigate the information captured by DrugCLIP’s learned pocket and ligand representations. A total of 335, 6690, 4967 and 5313 pockets-ligand pairs were encoded for COACH420, HOLO4K, scPDB and PDBBind, respectively.

First, the robustness of DrugCLIP’s molecule encoder to conformational variability was assessed by calculating the cosine similarity between embeddings of different conformations of the same molecule. A total of 716,413 pairs were evaluated across 1461 unique ligands represented at least twice (one pair) across the four data sets. The mean embedding similarity was calculated for each ligand, and the overall mean was obtained by averaging these ligand-level means, giving equal weight to each ligand regardless of the number of representative pairs.

Next, the relationship between molecule embedding similarity and chemical similarity was investigated. Canonical SMILES representations were retrieved from the CCD and extended connectivity fingerprints (ECFP4) and MACCS keys (166 predefined structural features) [124] were generated using RDKit. Chemical similarity was calculated using the Tanimoto coefficient for both fingerprint representations. Additionally, physicochemical similarity was assessed using ten molecular descriptors calculated by RDKit: molecular weight, logP, topological polar surface area (TPSA), hydrogen bond donor and acceptor counts, rotatable bond count, aromatic ring count, fraction of sp^3^ carbons, ring count, and heavy-atom count [125]. Descriptor values were standardised (z-scored) across the ligand set, and pairwise physicochemical similarity was calculated as 1 / (1 + Euclidean distance). The relationships between embedding similarity and each of the three chemical similarity measures were evaluated across 17,567,628 unique pairwise comparisons between 5928 distinct ligands using Spearman’s rank correlation coefficient (ρ) [126]. For ligands represented multiple times across the data sets, embedding similarity was averaged across all corresponding pairs of embeddings.

To further examine the relationship between ligand embedding similarity and chemical similarity, the mean cosine similarity between all pairs of pockets binding each ligand in a pair was also calculated. Ligand pairs were then stratified into seven intervals based on their mean pocket embedding cosine similarity, ranging from −0.75 to 1.00 in increments of 0.25. Although cosine similarity ranges from −1 to 1, no ligand pairs presented a mean pocket embedding similarity below −0.75. Within each interval, Spearman correlations were calculated between mean ligand embedding cosine similarity and ECFP4 Tanimoto similarity. Additionally, Spearman correlations were calculated between mean pocket and ligand embedding cosine similarities, and between mean pocket embedding cosine similarity and ligand ECFP4 Tanimoto similarity.

Finally, the link between pocket embedding and structural similarities was inspected. To obtain a suitable set of structurally superposable binding sites, comparisons were restricted to pockets binding the same ligand. This yielded 11,121 pockets binding to 1479 different ligands, resulting in 716,099 same-ligand pocket pairs across the four data sets. Ligand atoms were aligned using the SVDSuperimposer Biopython package [127], which implements the original algorithm by Kabsch [128] to determine the optimal rotation and translation for aligning two sets of corresponding points. The resulting transformations were applied to the protein atoms and pocket structural similarity was quantified using element-matched nearest-neighbour RMSD [129], similarly to Hoffmann *et al.* [130]. A high-confidence subset of 147,791 pocket pairs, comprising 9643 pockets and 1196 ligands was defined by retaining those pocket pairs with ligand RMSD ≤ 1 Å, hereafter referred to as “well-aligned”. Ten highly prevalent ligands, e.g., FAD, NAD, ADP, FMN or NAP, accounted for 89% of all comparisons. To avoid the overrepresentation of these frequently occurrent ligands, each ligand contributed a single observation, calculated as the mean pocket embedding cosine similarity and mean pocket RMSD across all corresponding same-ligand pairs.

### Bound ligand recovery benchmark

Jia *et al.* [36] evaluated DrugCLIP’s virtual screening performance using the Directory of Useful Decoys [80, 81] – Enhanced (DUD-E) [82] and Literature-derived PubChem BioAssay (LIT-PCBA) [83] benchmark data sets. Here, we assess DrugCLIP on a related task, referred to as “bound ligand recovery”. Given an experimentally determined protein-ligand complex, and a background library of candidate molecules, DrugCLIP was tasked with identifying the experimentally observed bound ligand among the candidates. Query pocket-ligand complexes were obtained from the subsets of COACH420, HOLO4K, scPDB and PDBBind employed by Sestak *et al.* [75]. The screening library (of presumed decoys, or negatives) consisted of on 48,999 non-ion ligands (>1 heavy atom) from the CCD [86], selected from the 50,022 ligands present in the PDBe [131, 132]. For each query, the experimentally bound ligand was considered the sole positive, with all remaining library molecules treated as assumed negatives.

A total of 17,191 pocket-ligand pairs were encoded by DrugCLIP across COACH420 (330), HOLO4K (6599), scPDB (4957) and PDBBind (5305). Additionally, the 48,999 ideal CCD ligand conformers were embedded. The entire CCD library was screened against each query pocket, and the rank of the experimentally bound ligand was recorded. Ideally, the bound ligand would rank first (#1), indicating that DrugCLIP assigns it a higher similarity score than all other compounds in the library. Performance was evaluated using three different metrics: top 1 recall or accuracy (%), enrichment factor (EF) at 1% (EF_1%_) and median rank. Top 1 recall measures the percentage of query pockets for which the experimentally bound ligand ranks ahead of all other 48,998 compounds in the CCD library. This represents the strictest evaluation criterion, for which relatively low values are expected. EF_1%_ measures the enrichment of active compounds within the top 1% of the ranked library relative to the random expectation. In our setting, where each query has a single positive (the experimentally bound ligand), EF_1%_ reflects how frequently the bound ligand is retrieved among the top 490 compounds (*_x_* = 0.01; *_M_* = 48,999) – (Equation 1). Under random ranking, the bound ligand would be expected to fall within this fraction for 1% of queries. For example, if the bound ligand ranked within the top 1% for 50 out of 100 query pockets, the corresponding recall would be 50% (Equations 2-3), yielding an EF_1%_ = 50, or a 50-fold enrichment over random expectation (Equation 4). Uncertainty in EF_1%_ and top 1 recall was quantified using 95% Wilson score confidence intervals [133]. Finally, the median rank of the bound ligand across all query pockets was also reported.

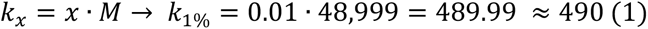

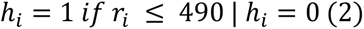

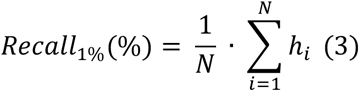

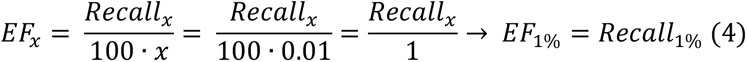

DrugCLIP’s pocket encoder was fine-tuned on approximately 40,000 protein-ligand complexes from BioLiP [87, 134], some of which may overlap with or resemble those in our benchmark data sets. To account for potential training-test overlap, a multi-level, multi-tiered leakage analysis was conducted. Pockets from the four data sets were pooled (*N* = 17,191) and deduplicated by PDB identifier and ligand code, yielding 15,533 unique pockets. This strategy was adopted to ensure sufficient sample size across the different leakage tiers. The first level examines protein structure and sequence similarity. The initial tier includes structures also present in DrugCLIP’s fine-tuning set, while subsequent tiers are defined by decreasing sequence identity (SI %) to the BioLiP fine-tuning set: >70, (50, 70], (30, 50] and ≤30. Pairwise sequence alignments and identity calculations were performed using MMseqs [110–112]. The second level examines ligand frequency, identity and chemical similarity, considering only complexes remaining after excluding overlapping PDBs and proteins sharing >30% SI with the (BioLiP) fine-tuning set. The initial tiers group ligands according to their frequency in the fine-tuning set: ≥100, 25-99, 5-24, 1-4 and 0 occurrences, the latter corresponding to ligands absent from the BioLiP set employed to fine-tune the model. For these unseen ligands, chemical similarity to molecules in the BioLiP fine-tuning set was assessed using the Tanimoto coefficient, with the following intervals: [0.9, 1.0], [0.7, 0.9), [0.5, 0.7), [0.3, 0.5) and <0.3 (novel chemistry). This sequential and cumulative filtering strategy enables the evaluation of DrugCLIP’s performance across progressively stricter levels of training-test separation, providing insight into its potential reliance on memorisation and its ability to generalise to previously unseen proteins and ligands.

It is important to note that this leakage analysis is restricted to the BioLiP complexes used to fine-tune DrugCLIP. Its molecular encoder was initialised from Uni-Mol [135], which was pretrained on molecular structures, whereas its pocket encoder was pretrained using the protein fragment-surrounding alignment (ProFSA) framework [36], involving approximately 5.5 million synthetic pseudo-pocket-ligand pairs. Although prior exposure to individual molecules or protein structures during these earlier stages cannot be excluded, such exposure does not necessarily imply prior knowledge of the experimentally observed protein-ligand binding relationships evaluated here. Therefore, our analysis assesses generalisation beyond the fine-tuning set, rather than complete independence from all pretraining data.

### Performance robustness and degradation analysis

DrugCLIP’s performance across tasks, including ligand recovery, virtual screening and pocket matching, relies on the cosine similarity between the learned representations generated by its pocket and molecule encoders. The bound ligand recovery benchmark described above, represents an ideal scenario, in which binding pockets are defined from an experimentally determined protein-ligand complex, with sidechain atoms already oriented to accommodate the observed ligand. However, this is not necessarily the case in real-world virtual screening applications, where an experimental structure of the target protein might me unavailable, only alternative *holo* or *apo* conformations may be accessible or experimentally determined ligand binding information could be lacking, requiring binding pocket prediction as a starting point. To investigate the impact of these factors on DrugCLIP’s performance, several experiments were carried out in which either the pocket or ligand was modified and re-encoded to assess the resulting changes in pocket-ligand similarity scores and derived retrieval performance. These included using the experimentally bound ligand conformation instead of the generic CCD conformer, replacing experimentally defined pockets with predicted ones, and using predicted structure models instead of experimental structures.

Three-dimensional models predicted by AlphaFold [136–138] were downloaded from the AlphaFold Protein Structure Database (AFDB) [139–142] for 4854 unique entries in UniProt [143, 144], corresponding to 12,930 of the 15,533 unique pockets pooled across the four data sets. Pockets were transferred from the experimental PDB structures to the corresponding AFDB models using chain and residue mappings between PDBe and UniProt obtained through SIFTS [145, 146]. Thus, pockets in the predicted structures were defined using the same set of residues located within 6 Å of the bound ligand in the experimental structure. These transferred pockets were encoded using DrugCLIP, and their embeddings compared with those of the corresponding PDB pockets. This analysis aimed to assess the impact of using predicted rather than experimentally determined protein structures on DrugCLIP’s pocket representations and performance.

The effects of protein flexibility and alternative (apo/holo) pocket conformations on DrugCLIP’s performance were assessed using the Apo-Holo Juxtaposition Database (AHoJ-DB) [147, 148]. AHoJ-DB (https://apoholo.cz/db) provides precomputed apo-holo structure matches for biologically relevant ligands from BioLiP [134]. Each entry corresponds to a target PDB chain and a bound ligand and contains a list of matching structures classified as “HOLO” (bound) or “APO” (unbound) relative to the binding site defined by the ligand. Information from 162,454 apo and 1,032,881 holo structures was retrieved across the four data sets, corresponding to 5244 and 9620 unique pockets with at least one apo and holo conformation, respectively. These large counts arise because AHoJ-DB operates at the level of individual pockets, rather than PDB entries or chains, aggregating matching chains across all PDB structures associated with the same UniProt accession and binding site. Consequently, a single pocket may have corresponding conformations in thousands of chains. Binding pockets in both apo and holo structures were transferred from the original ligand-bound complex using SIFTS mappings. Thus, pockets in alternative holo structures were defined by the residues interacting with the ligand in the original complex, rather than those interacting with the ligand in the matched holo structure. Similarly, apo pockets were defined using the same residues, despite the absence of the ligand in the unbound conformation. These alternative pocket conformations were encoded using DrugCLIP, and their embeddings compared with those of the corresponding original ligand-bound pockets. Pearson’s correlation coefficient (r) [149] was used to assess the relationship between the pocket-ligand cosine similarity scores obtained from the original pockets and their alternative apo/holo conformations.

DrugCLIP’s dependence on an accurate pocket definition and binding residue selection was further tested by replacing experimentally observed binding sites with predicted pockets. P2Rank [72, 150] was used to predict ligand binding sites in both the original experimental PDB structures and their corresponding AFDB models. Predicted pockets matching experimentally observed sites were encoded, and their pocket-ligand cosine similarity scores compared with those from the experimentally defined pockets. Predictions that did not match any observed binding site (false positives) were also encoded and analysed as a negative control. For these pockets, a distribution of cosine similarity scores centred around 0 would be expected, reflecting the absence of a known binding relationship with the ligand under evaluation. Further details of the pocket prediction procedure are provided in the following Methods subsection.

In addition, DrugCLIP’s robustness to variations in ligand conformation was further assessed. For all 15,533 unique pocket-ligand pairs pooled across the four data sets, the experimentally observed bound ligand conformation was encoded using DrugCLIP, and its cosine similarity score against the corresponding pocket embedding was compared with that obtained using the ideal ligand conformer from the CCD.

Finally, three types of negative controls were evaluated: incorrect P2Rank pocket predictions from experimental PDB structures, incorrect P2Rank predictions from AlphaFold models, and randomly generated protein surface patches. Incorrect predictions were defined as those not matching any experimentally observed binding site. Further details on the generation of random surface patches are provided in the following Methods subsection.

### Pocket prediction

Numerous methods for predicting ligand binding sites have been developed over the past three decades, reflecting substantial advances and shifts in methodological approaches [91, 151–158]. P2Rank [159–161] has demonstrated strong predictive performance, as shown in the recent benchmark on the LIGYSIS data set [6, 162] by Utgés and Barton [74]. Combined with its computational efficiency and support for parallel execution, this makes it particularly suitable for large-scale pocket prediction. Accordingly, P2Rank was used with its default configuration to predict binding pockets in both PDB structures and AFDB models throughout this work, without applying any additional filtering based on prediction scores.

Several criteria can be used to classify predicted pockets as correct or incorrect. The most widely used are based on the Euclidean distance (Å) between the predicted and observed pocket centroids (DCC), or between the predicted centroid and the nearest ligand heavy atom (DCA) and typically use a threshold of 4 Å [163–165]. Predictions satisfying either DCA or DCC ≤ 4 Å were considered correct. Using these criteria, pocket recall was 75.2% (248/330), 83.9% (5536/6599), 89.7% (4448/4957) and 87.1% (4623/5305) for COACH420, HOLO4K, scPDB and PDBBind, respectively, or 85.8% (13,327/15,533) for the pooled set of unique pockets – deduplicated by PDB ID and ligand code. To evaluate predictions on AFDB models, the experimental PDB structures, including their bound ligands, were structurally aligned to their corresponding predicted models. The transformed coordinates were then used to calculate DCA and DCC following the same procedure applied to the experimental structures. Pocket recall was lower for AFDB models, reaching 59.4% (148/249), 65.9% (2223/3374), 75.3% (2099/2788) and 64.4% (2334/3624) for COACH420, HOLO4K, scPDB, PDBBind, respectively, and 67.8% (6804/10,035) for the pooled set of unique pockets. The number of evaluated pockets was smaller for AFDB models than for experimental structures – 10,035 *vs* 15,533 – as only pockets with available AFDB models and complete one-to-one residue- and atom-level mappings between PDB structures and AFDB models were considered (64.6%).

Other approaches to evaluating predicted pockets rely on volume, shape, or residue overlap. Volume- or shape-based criteria include discretised volume overlap (DVO) [166], relative volume overlap (RVO) [74] and atom overlap (OVR) [120], among others. These metrics assess how closely the predicted pocket envelope matches the experimentally observed one and often require volumetric calculations that can be computationally expensive at scale. Residue overlap-based criteria include relative residue overlap (RRO) [74] and residue intersection over union (IOU), also known as Jaccard index (JI) [167]. These quantify the agreement in residue membership between predicted and observed pockets. Unlike distance-based criteria, there is no clear consensus in the literature regarding the thresholds used to classify predictions as correct based on volume or residue overlap. Consequently, these metrics are often used to assess pocket prediction quality after predictions have been classified as true positives using other criteria. In this work, we examine the relationships between pocket embedding similarity and two residue overlap metrics: IOU and recall (RRO).

### Random surface patch generation

Solvent accessible surface area (SASA) was calculated using the Biopython [127] implementation of the original “rolling ball” algorithm by Shrake and Rupley [168, 169]. SASA values were normalised to relative solvent accessibility (RSA) following the method of Tien *et al.* [170]. Surface residues were defined as those presenting RSA > 0.02 (2%). Residues lining the experimental ligand binding site (within 6 Å of the bound ligand) were excluded to ensure that no generated patch shared residues with the true binding site. The remaining surface residues were grouped into spatially adjacent regions or patches no larger than 128 heavy atoms. Finally, any groups presenting fewer than 40 heavy atoms were merged into nearby patches based on the Euclidean distance between their centroids, subject to a maximum of 511atoms per patch, corresponding to DrugCLIP’s default limit on the number of pocket atoms. In total, 252,222 unique surface patches were generated and encoded across the four benchmark data sets: COACH420 (3579), HOLO4K (64,414), scPDB (102,044) and PDBBind (82,185). Representative examples of these patches on protein surfaces are shown in Figure 7, with further details provided in Figure S5 and Table S7.

**Figure 7.**
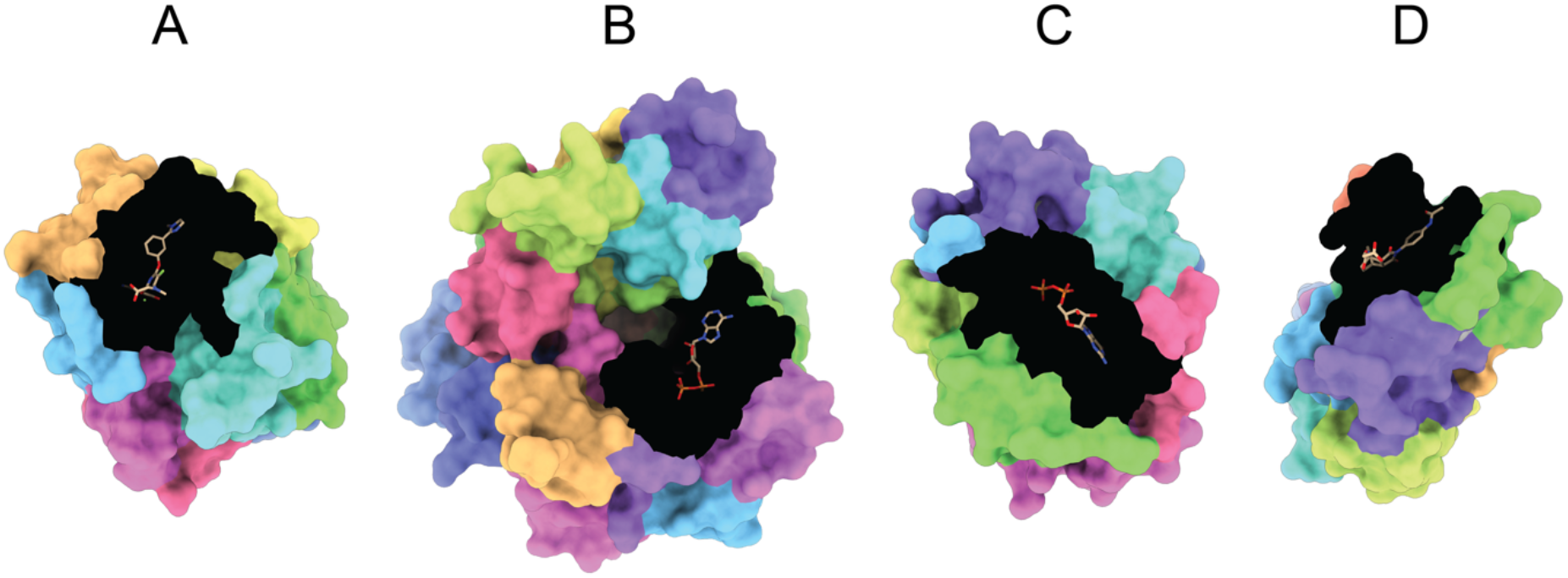
Random surface patches. Four representative examples of randomly generated protein surface patches used as negative controls for DrugCLIP. A total of 252,222 patches were generated from 13,723 unique PDB chains pooled across the four benchmark data sets: COACH420, HOLO4K, scPDB and PDBBind. In each panel, the experimentally observed binding pocket (residues within 6 Å of the bound ligand) is shown in black, with the ligand represented as sticks coloured by heteroatom. The remaining solvent-accessible protein surface is coloured to distinguish the individual patches. **(A)** Human coagulation factor X (P00742) bound to Fidexaban (Z34) – PDB: 1FJS [171]; **(B)** Chicken poly(ADP-ribose) polymerase 1 (P26446) bound to carbanicotinamide-adenine-dinucleotide (CNA) – PDB: 1A26 [172]; **(C)** Dog GTP-binding nuclear protein Ran (P62825) bound to guanosine-5’-diphosphate (GDP) – PDB: 1A2K [173]; **(D)** Mouse Fab antibody binding to [4-(4-acetyla-mino-phenyl)-3,5-dioxo-4-aza-tricyclo[5.2.2.02,6]-undec-1-ylcarbamoylo-xy]-acetic acid (FRA) – PDB: 1A4K [174]. Structure visualisation with ChimeraX [175–177].

### Statistics and Reproducibility

Data analysis was carried out primarily with the following Python libraries: NumPy [178], Pandas [179] and SciPy [180]. RDKit [109] and Biopython [127] were used for cheminformatics and structural bioinformatics processing, with Matplotlib [181] for plotting. All statistical tests performed are two-tailed, and significance level α = 0.05.

## List of Abbreviations

3D: three-dimensional.
ADP: adenosine-5’-diphosphate.
ATP: adenosine-5’-triphosphate.
AFDB: AlphaFold Protein Structure Database.
AHoJ-DB: Apo-Holo Juxtaposition Database.
AUROC: area under the receiver operating characteristic curve.
AUC: area under the curve.
CCD: Chemical Component Dictionary.
CI: confidence interval.
CLIP: contrastive language-image pretraining.
DCA: distance from predicted centroid to closest ligand atom.
DCC: distance from (predicted) centroid to (observed) centroid.
DUD: Directory of Useful Decoys.
DUD-E: Directory of Useful Decoys – Enhanced.
ECFP: extended connectivity fingerprint.
EF: enrichment factor.
EPoCS: ESM-based Pocket Cross Similarity.
FAD: flavin-adenine dinucleotide.
FMN: flavin mononucleotide.
ID: identifier.
IQR: interquartile range.
LIT-PCBA: Literature-derived PubChem BioAssay.
LLM: lead-like molecule.
LMDB: lightning memory-mapped database.
MACCS: Molecular Access System.
MOAD: Mother of all Databases.
NAD: nicotinamide-adenine-dinucleotide.
NAP: nicotinamide-adenine-dinucleotide phosphate.
NMR: nuclear magnetic resonance.
PDB: Protein Data Bank.
PDBe: Protein Data Bank Europe.
ProFSA: protein fragment-surrounding alignment.
ProSPECCTs: Protein Site Pairs for the Evaluation of Cavity Comparison Tools.
PSI: pocket sequence identity.
RMSD: root mean square deviation.
ROC: receiver operating characteristic curve.
RSA: relative solvent accessibility.
SASA: solvent accessible surface area.
SI: sequence identity.
SIFTS: Structure Integration with Function, Taxonomy and Sequence.
SMILES: simplified molecular input line entry system.
TC: Tanimoto coefficient.
TPSA: topological polar surface area.
TS: Tanimoto similarity.

## Data and Software Availability

The main results, tables and files necessary to replicate the analysis described in this work can be found on Zenodo: https://zenodo.org/records/23812210 [182, 183]. The software developed to carry out the analyses described in this work can be found on GitHub: https://github.com/JavierSanchez-Utges/DrugCLIP-pocket-analysis [184, 185].

## Author Contributions

J.S.U. and C.O. conceived, designed and developed the research. J.S.U. wrote the software and analysed the data. J.S.U. wrote the original manuscript. J.S.U. and D.T.J. reviewed and edited the manuscript. C.O. secured funding and supervised.

## Acknowledgements

We thank the rest of the Orengo (CATH) group, in particular David Wagemann, as well as Jude Wells, and Drs Radoslav Krivák and Christos Feidakis, for their comments and insightful discussions about this project. We extend our gratitude to the computational sciences (CS) IT department for their support of the HPC infrastructure this study was carried out on, particularly Angie Chan, Ed Gair and Jamie O’Connor.

## Supporting Information

Additional ProSPECCTs pocket matching benchmark results and comparisons with previously published methods; computational runtime comparison between DrugCLIP and PocketVec; analysis of ligand embedding similarity in relation to chemical and binding pocket similarity; evaluation of predicted pocket quality and its influence on virtual screening performance; pairwise comparisons of ligand retrieval performance across screening conditions; relationships between pocket-ligand similarity scores and ligand retrieval ranks; sample sizes and enrichment factors across de-leakage levels; and characterisation of random protein surface patches used as negative controls.

