## Supplementary Material for "Interrogating contrastive learning embeddings for structure-based virtual screening: a case study on DrugCLIP"

### Supporting Information

**Javier S. Utgés, David T. Jones, Christine Orengo***

Institute of Structural and Molecular Biology, Darwin Building, University College London, Gower Street, London, WC1E 6BT, UK.

| Method | Mean AUROC |
| --- | --- |
| DrugCLIP | **0.86** |
| DrugCLIP$_{12\ Å}$ | 0.76 |
| DrugCLIP$_{Sim\ Vec}$ | 0.78 |
| DrugCLIP$_{Rank\ Vec}$ | 0.81 |
| EPoCS | 0.78 |
| NRGRank | 0.61 |
| PocketVec | 0.74 |
| Sequence | 0.65 |
| Ligand Code | 0.66 |
| PSI | 0.80 |
| ECFP4 TS | 0.84 |

**Table S1. Mean AUROC across ProSPECCTs pocket matching benchmark data sets.** DrugCLIP [1], EPoCS [2] and PocketVec [3] methods are default. NRGRank [4] is used to screen the same 128 compounds employed by PocketVec and generate a pocket descriptor in the same fashion. Three DrugCLIP variants were also tested: one that defines pockets using a 12 Å threshold (DrugCLIP$_{12\ Å}$) – as opposed the default 6 Å threshold – from the bound ligand atoms, one that uses DrugCLIP's screening ability to generate a pocket-ligand similarity vector using the PocketVec 128 LLMs (DrugCLIP$_{Sim\ Vec}$), and DrugCLIP$_{Rank\ Vec}$, which ranks the 128 molecules by their screening score. Four baselines are also included for reference, which use a sequence hash, or 100% sequence identity (Sequence), the ligand code (Ligand code), pocket sequence identity (PSI) or Tanimoto similarity between bound ligands (ECFP4 TS) to assign classification labels.

| | **D1** | **D1.2** | **D2** | **D3** | **D4** | **D5** | **D5.2** | **D6** | **D7** |
|---|---|---|---|---|---|---|---|---|---|
| DrugCLIP | 0.95 | **1.00** | **1.00** | 0.66 | 0.65 | **0.93** | **0.92** | **0.84** | **0.92** |
| $\text{DrugCLIP}_{12\,\text{Å}}$ | 0.98 | **1.00** | 0.99 | 0.73 | 0.68 | 0.69 | 0.67 | 0.30 | 0.91 |
| $\text{DrugCLIP}_{\text{Sim Vec}}$ | 0.84 | 0.97 | 0.98 | 0.66 | 0.61 | 0.79 | 0.79 | 0.71 | 0.82 |
| $\text{DrugCLIP}_{\text{Rank Vec}}$ | 0.89 | 0.99 | 0.98 | 0.71 | 0.65 | 0.85 | 0.84 | 0.74 | 0.84 |
| EPoCS | **1.00** | **1.00** | **1.00** | 0.82 | 0.84 | 0.59 | 0.56 | 0.51 | 0.86 |
| NRGRank | 0.73 | 0.83 | 0.62 | 0.63 | 0.55 | 0.56 | 0.54 | 0.55 | 0.79 |
| PocketVec | 0.97 | **1.00** | 0.96 | 0.67 | 0.72 | 0.64 | 0.62 | 0.44 | 0.87 |
| SiteContext* | 0.92 | **1.00** | 0.98 | 0.55 | 0.57 | 0.65 | 0.75 | 0.53 | 0.81 |
| SiteMine* | **1.00** | – | **1.00** | 0.69 | 0.72 | 0.78 | – | 0.61 | 0.91 |
| BindSiteS-CNN* | 0.94 | 0.98 | 0.83 | 0.91 | 0.79 | 0.64 | 0.66 | 0.62 | 0.78 |
| Site2Vec* | **1.00** | 0.94 | **1.00** | **0.99** | **0.99** | – | 0.86 | 0.53 | 0.66 |
| DeeplyTough* | 0.95 | 0.98 | 0.90 | 0.76 | 0.75 | 0.67 | 0.63 | 0.54 | 0.83 |
| IsoMIF† | 0.77 | 0.97 | 0.70 | 0.59 | 0.59 | 0.75 | 0.81 | 0.62 | 0.87 |
| RAPMAD† | 0.85 | 0.83 | 0.82 | 0.61 | 0.63 | 0.55 | 0.52 | 0.60 | 0.74 |
| SiteHopper† | 0.98 | 0.94 | **1.00** | 0.75 | 0.75 | 0.72 | 0.81 | 0.56 | 0.77 |
| KRIPO† | 0.91 | **1.00** | 0.96 | 0.60 | 0.61 | 0.76 | 0.77 | 0.73 | 0.85 |
| TIFP† | 0.66 | 0.90 | 0.91 | 0.66 | 0.66 | 0.71 | 0.63 | 0.55 | 0.71 |
| Grim† | 0.69 | 0.97 | 0.92 | 0.55 | 0.56 | 0.69 | 0.61 | 0.45 | 0.70 |
| Volsite/Shaper† | 0.93 | 0.99 | 0.78 | 0.68 | 0.76 | 0.56 | 0.58 | 0.71 | 0.77 |
| Shaper† | 0.96 | 0.93 | 0.93 | 0.71 | 0.76 | 0.65 | 0.65 | 0.54 | 0.75 |
| FuzCav† | 0.94 | 0.99 | 0.99 | 0.69 | 0.58 | 0.55 | 0.54 | 0.67 | 0.77 |
| ProBiS† | **1.00** | **1.00** | **1.00** | 0.47 | 0.46 | 0.54 | 0.55 | 0.50 | 0.85 |
| SMAP† | **1.00** | **1.00** | **1.00** | 0.76 | 0.65 | 0.62 | 0.54 | 0.68 | 0.86 |
| PocketMatch† | 0.82 | 0.98 | 0.96 | 0.59 | 0.57 | 0.66 | 0.60 | 0.51 | 0.82 |
| SiteAlign† | 0.97 | **1.00** | **1.00** | 0.85 | 0.80 | 0.59 | 0.57 | 0.44 | 0.87 |
| TM-align† | **1.00** | **1.00** | **1.00** | 0.49 | 0.49 | 0.66 | 0.62 | 0.59 | 0.88 |
| SiteEngine† | 0.96 | **1.00** | **1.00** | 0.82 | 0.79 | 0.64 | 0.57 | 0.55 | 0.86 |
| Cavbase† | 0.98 | 0.91 | 0.87 | 0.65 | 0.64 | 0.60 | 0.57 | 0.55 | 0.82 |

| | | | | | | | | | |
|---|---|---|---|---|---|---|---|---|---|
| Sequence | 0.67 | 0.89 | 1.00 | 0.67 | 0.67 | 0.54 | 0.54 | 0.50 | 0.71 |
| Ligand Code | 0.51 | 0.59 | 0.99 | 0.51 | 0.51 | 0.97 | 0.95 | 0.47 | 0.77 |
| PSI | **1.00** | **1.00** | **1.00** | **1.00** | **1.00** | 0.57 | 0.55 | 0.41 | 0.86 |
| ECFP4 TS | 0.79 | **1.00** | **1.00** | 0.83 | 0.83 | 0.97 | 0.94 | 0.46 | 0.85 |

**Table S2. Extended ProSPECCTs pocket matching benchmark.** Extension of the ProSPECCTs pocket matching benchmark presented in the main text. All reported values correspond to AUROC. In addition to the seven methods [1-4] and four baselines included in the main evaluation, an additional 21 methods are now considered for further reference. These comprise 16 methods evaluated by Ehrt *et al.* [5] in 2018 – Cavbase [6], SiteEngine [7], TM-align [8], SiteAlign [9], PocketMatch [10], SMAP [11], ProBiS [12], FuzCav [13], Volsite/Shaper and Shaper [14], Grim [15], KRIPO [16], TIFP [15], SiteHopper [17], RAPMAD [18] and IsoMIF [19] – as well as five more recent methods: DeeplyTough [20], Site2Vec [21], BindSiteS-CNN [22], SiteMine [23] and SiteContext [24]. Superscripts identify results obtained from previously published studies rather than generated in this work. Specifically, a dagger (†) denotes results reported by Ehrt *et al.* [5] in their 2018 assessment, while an asterisk (*) indicates results taken from the corresponding method's original publication. AUROC values may differ slightly from those reported in the main manuscript, as the values presented here were calculated using the full (redundant) ProSPECCTs pocket-pair sets to ensure consistency with previous studies. The highest AUROC for each data set is highlighted in bold. Dashes (–) indicate that results are unavailable for a given method on a particular ProSPECCTs set.

| Method | Mean AUROC | # sets included | Missing sets |
|---|---|---|---|
| DrugCLIP | **0.87** | 9 | – |
| DrugCLIP$_{12\,Å}$ | 0.77 | 9 | – |
| DrugCLIP$_{Sim\ Vec}$ | 0.80 | 9 | – |
| DrugCLIP$_{Rank\ Vec}$ | 0.83 | 9 | – |
| EPoCS | 0.80 | 9 | – |
| NRGRank | 0.64 | 9 | – |
| PocketVec | 0.77 | 9 | – |
| SiteContext* | 0.75 | 9 | – |
| SiteMine* | *0.82* | 7 | D1.2, D5.2 |
| BindSiteS-CNN* | 0.79 | 9 | – |
| Site2Vec* | ***0.87*** | 8 | D5 |
| DeeplyTough* | 0.78 | 9 | – |
| IsoMIF† | 0.74 | 9 | – |
| RAPMAD† | 0.68 | 9 | – |
| SiteHopper† | 0.81 | 9 | – |
| KRIPO† | 0.80 | 9 | – |
| TIFP† | 0.71 | 9 | – |
| Grim† | 0.68 | 9 | – |
| Volsite/Shaper† | 0.75 | 9 | – |
| Shaper† | 0.76 | 9 | – |
| FuzCav† | 0.75 | 9 | – |
| ProBiS† | 0.71 | 9 | – |
| SMAP† | 0.79 | 9 | – |
| PocketMatch† | 0.72 | 9 | – |
| SiteAlign† | 0.79 | 9 | – |
| TM-align† | 0.75 | 9 | – |
| SiteEngine† | 0.80 | 9 | – |
| Cavbase† | 0.73 | 9 | – |
| Sequence | 0.69 | 9 | – |

| | | | |
|---|---|---|---|
| Ligand Code | 0.70 | 9 | – |
| PSI | 0.82 | 9 | – |
| ECFP4 TS | 0.85 | 9 | – |

**Table S3. Mean AUROC across ProSPECCTs pocket matching benchmark sets (extension).** Superscripts identify results obtained from previously published studies rather than generated in this work. A dagger (†) denotes results reported by Ehrt *et al.* [5] in their 2018 assessment, while an asterisk (*) indicates results taken from the corresponding method's original publication. The method(s) achieving the highest mean AUROC are highlighted in bold, while italics indicate that the mean was calculated using fewer than the nine ProSPECCTs sets due to missing results.

| Attribute | PocketVec | DrugCLIP |
|---|---|---|
| Hardware | Intel Xeon E3-1240 v2/v3 (3.4 GHz), Gold 5118 (2.3 GHz), E5-2650 v3 (2.3 GHz) | NVIDIA A40 (46 GB) |
| Parallelisation | ≤ 20 concurrent single-core jobs | Single GPU |
| Time per pocket | 80.0 min (mean) | ≤ 0.15 s (16.7/115) |
| Total compute | 153.3 CPU-hours | 16.7 GPU-seconds |
| Wall-clock time | 11.2 h | 16.7 s |
| Speed-up | 1× | $10^4\times$ |

**Table S4. Pocket encoding runtime comparison between PocketVec and DrugCLIP.** This analysis of the computational cost was carried out on the D6 subset of the ProSPECCTs suite, also referred to as the Barelier set [25], which includes 115 pockets. For PocketVec, the workload per pocket entails cavity definition, rDock [26] docking of 128 lead-like molecules (× 25 runs) and descriptor generation. For DrugCLIP, the timing includes Python start-up, model build, loading of the 6 folds' checkpoints and pocket encoding – one forward pass through each of the six model folds (batch size 32). Pre-processing is excluded for both methods. This includes receptor MOL2 and pocket centroid generation for PocketVec and pocket extraction and LMDB construction for DrugCLIP. Pocket similarity is calculated as the cosine similarity between the vectors generated by the two methods. DrugCLIP is on average four orders of magnitude (at least 10,000) faster than PocketVec.

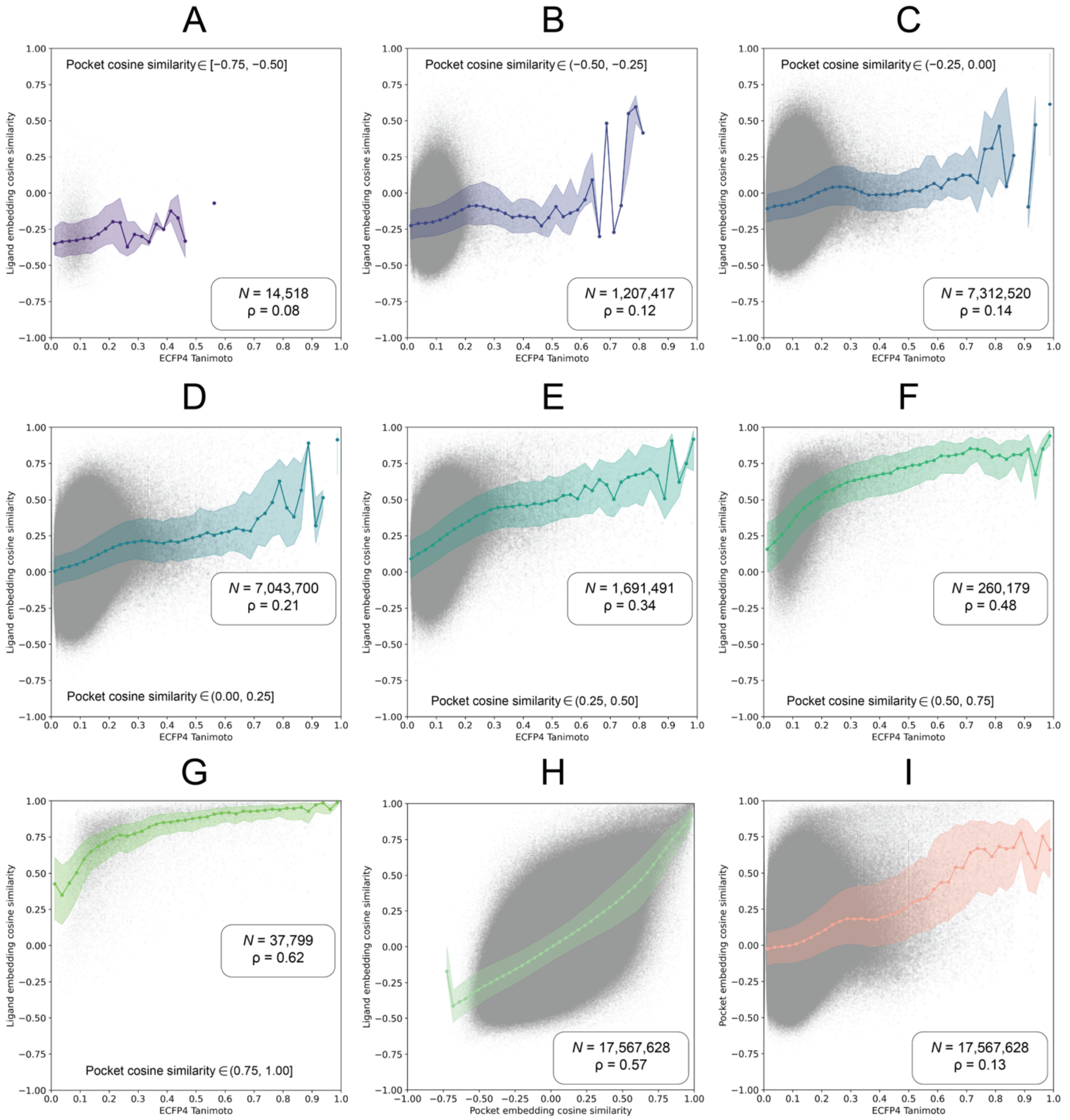


**Figure S1. Relationship between DrugCLIP ligand embedding cosine similarity and chemical similarity stratified by binding pocket similarity. (A-G)** Spearman correlations between ligand embedding cosine similarity and ECFP4 Tanimoto similarity for ligand pairs stratified by the cosine similarity of their corresponding pocket embeddings, across seven intervals: **(A)** −0.75 to −0.50; **(B)** −0.50 to −0.25; **(C)** −0.25 to 0.00; **(D)** 0.00 to 0.25; **(E)** 0.25 to 0.50; **(F)** 0.50 to 0.75; **(G)** 0.75 to 1.00; **(H)** Relationship between pocket and ligand embedding cosine similarities; **(I)** Relationship between pocket embedding cosine similarity and ligand ECFP4 Tanimoto similarity. Grey points represent individual (ligand) pairwise comparisons, coloured lines indicate binned median values, and shaded regions represent variability within bins (IQR). The number of pairwise comparisons ($N$) and Spearman correlation coefficient ($\rho$) are indicated in each panel.

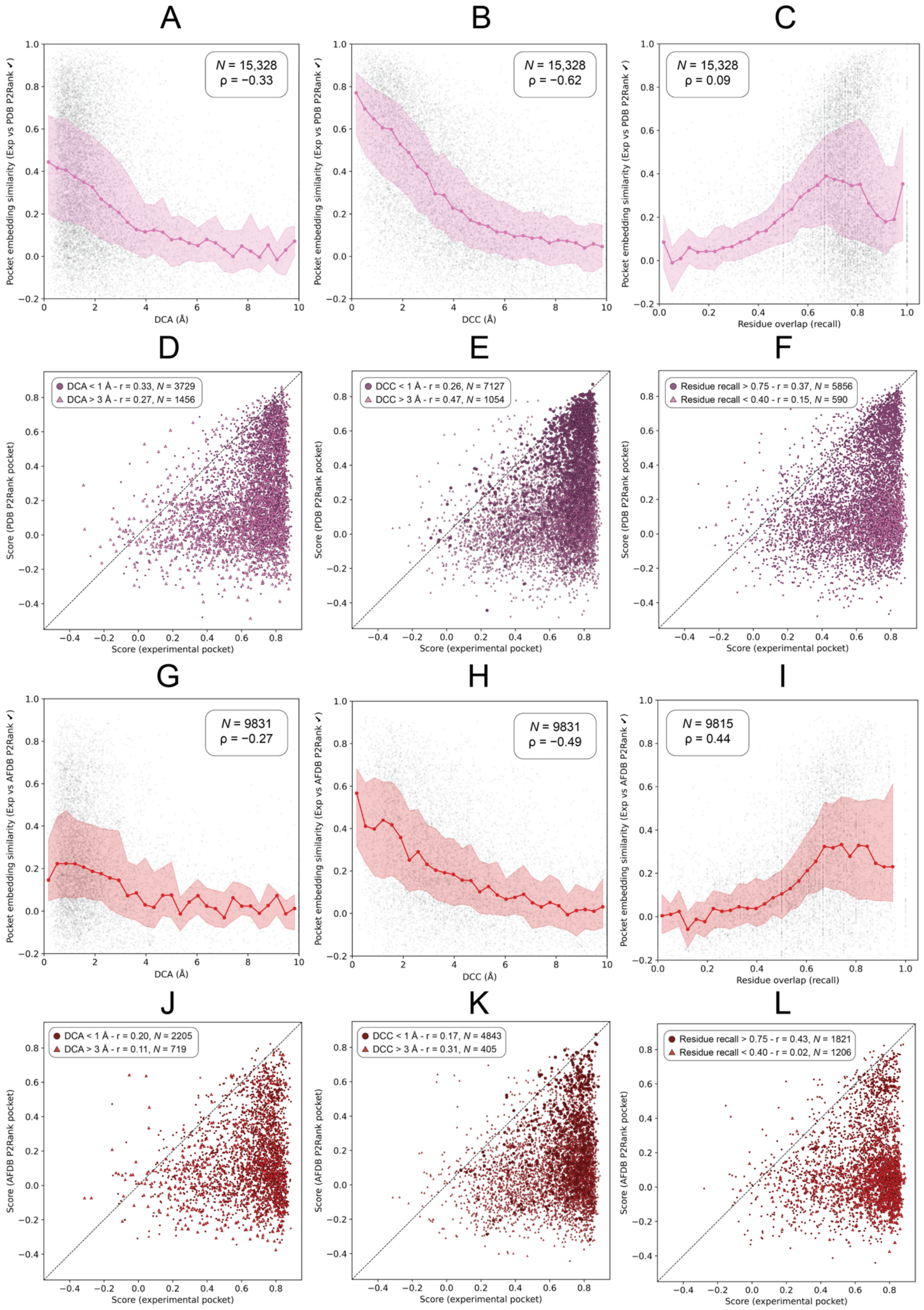


**Figure S2. Why does DrugCLIP perform worse on predicted pockets?** These panels complement Figure 5 by examining the relationships between experimental and predicted pocket embedding similarities, their respective pocket-ligand scores, and three other criteria used to classify pocket predictions as correct or incorrect: DCA, DCC and residue recall (relative to the observed pocket). Pearson's correlation was used to assess linear relationships between screening scores, whereas Spearman's rank correlation was employed to evaluate monotonic associations involving residue overlap, spatial distances and embedding similarities. **(A-C, G-I)** Scatter plots illustrating these relationships for DCA, DCC and residue recall, respectively, with trends represented by the median and interquartile range (IQR) within X-axis bin; **(D-F, J-L)** Relationships between experimental (X-axis) and predicted (Y-axis) pocket-ligand scores, with points coloured according to their corresponding pocket selection criteria. Sample sizes (*N*) and correlation coefficients (r, ρ) are indicated for reference.

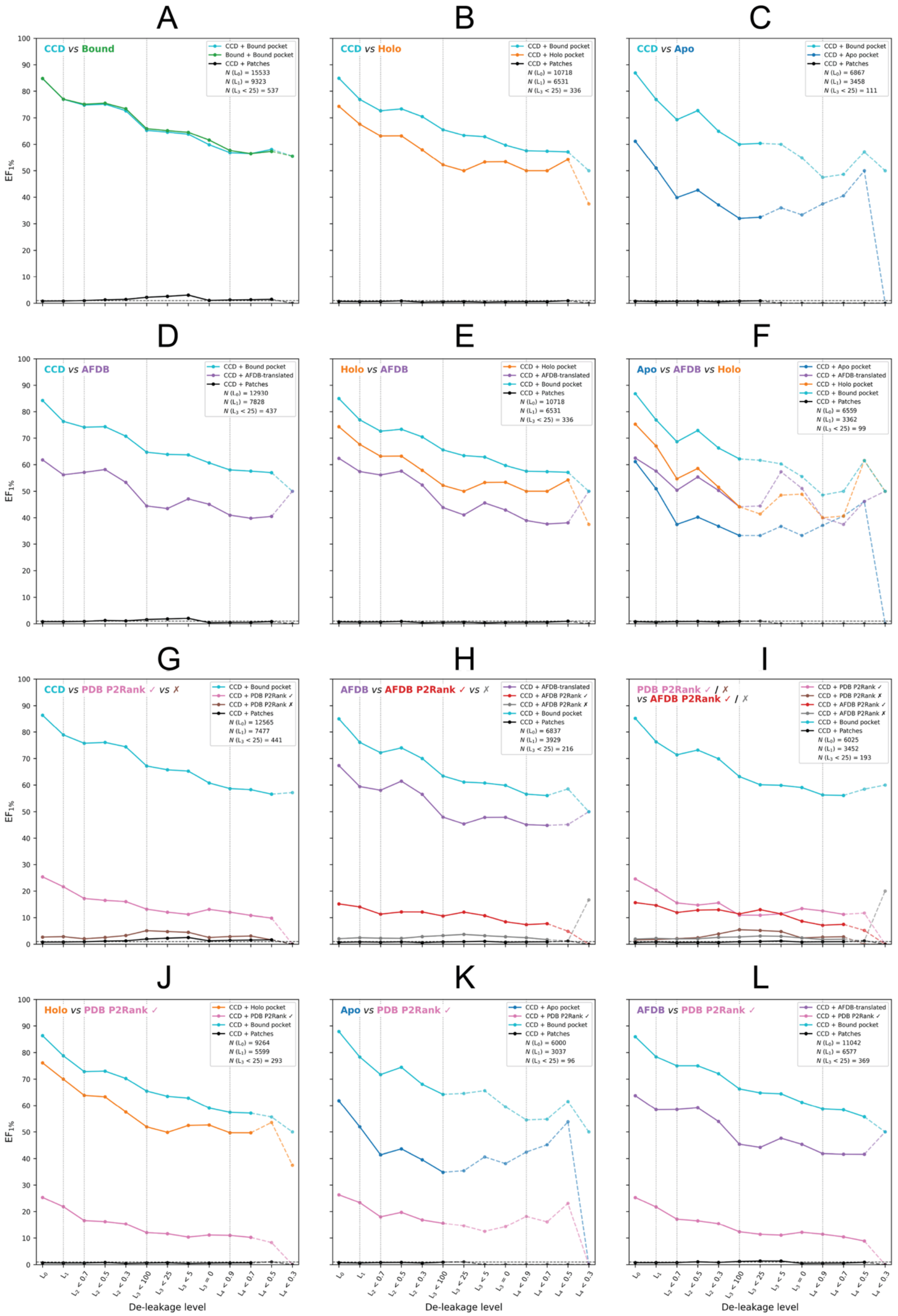


**Figure S3. Pairwise comparison of DrugCLIP ligand retrieval performance across different experimental conditions.** Enrichment factor at the top 1% ($EF_{1\%}$) of the screening library is shown across the 13 levels of the de-leakage ladder for different pocket and ligand representations evaluated on matched subsets of ligand binding pockets. **(A)** Generic CCD [27] ligand conformer (default) *vs* bound ligand conformer against the crystal pocket; **(B)** CCD *vs* alternative holo pocket; **(C)** CCD *vs* apo pocket conformation; **(D)** CCD *vs* AFDB-translated pocket (same residues); **(E)** Holo *vs* AFDB-translated; **(F)** Apo *vs* AFDB-translated *vs* holo; **(G)** Correctly and incorrectly predicted P2Rank pockets on experimental structures; **(H)** AFDB-translated *vs* correctly and incorrectly predicted P2Rank [28] pockets on AFDB [29] structures; **(I)** Correctly and incorrectly predicted P2Rank pockets on experimental and AFDB models; **(J)** Holo *vs* correctly predicted P2Rank pockets on experimental structures – (holo) conformational error *vs* pocket prediction error; **(K)** Apo *vs* correctly predicted P2Rank pockets on experimental structures – (apo) conformational error *vs* pocket prediction error; **(L)** AFDB-translated *vs* correctly predicted P2Rank pockets on experimental structures – structure prediction error *vs* pocket prediction error. CCD ligand conformer *vs* experimental pocket and random surface patches are included as positive and negative controls, respectively, where applicable. All comparisons within each panel are restricted to the same subset of ligand binding sites to ensure direct comparability between conditions. Solid lines represent de-leakage levels with at least 100 ligand-binding sites, while dashed lines indicate levels with fewer than 100. The number of pockets evaluated at $L_0$, $L_1$ and $L_3 < 25$ are indicated in each panel.

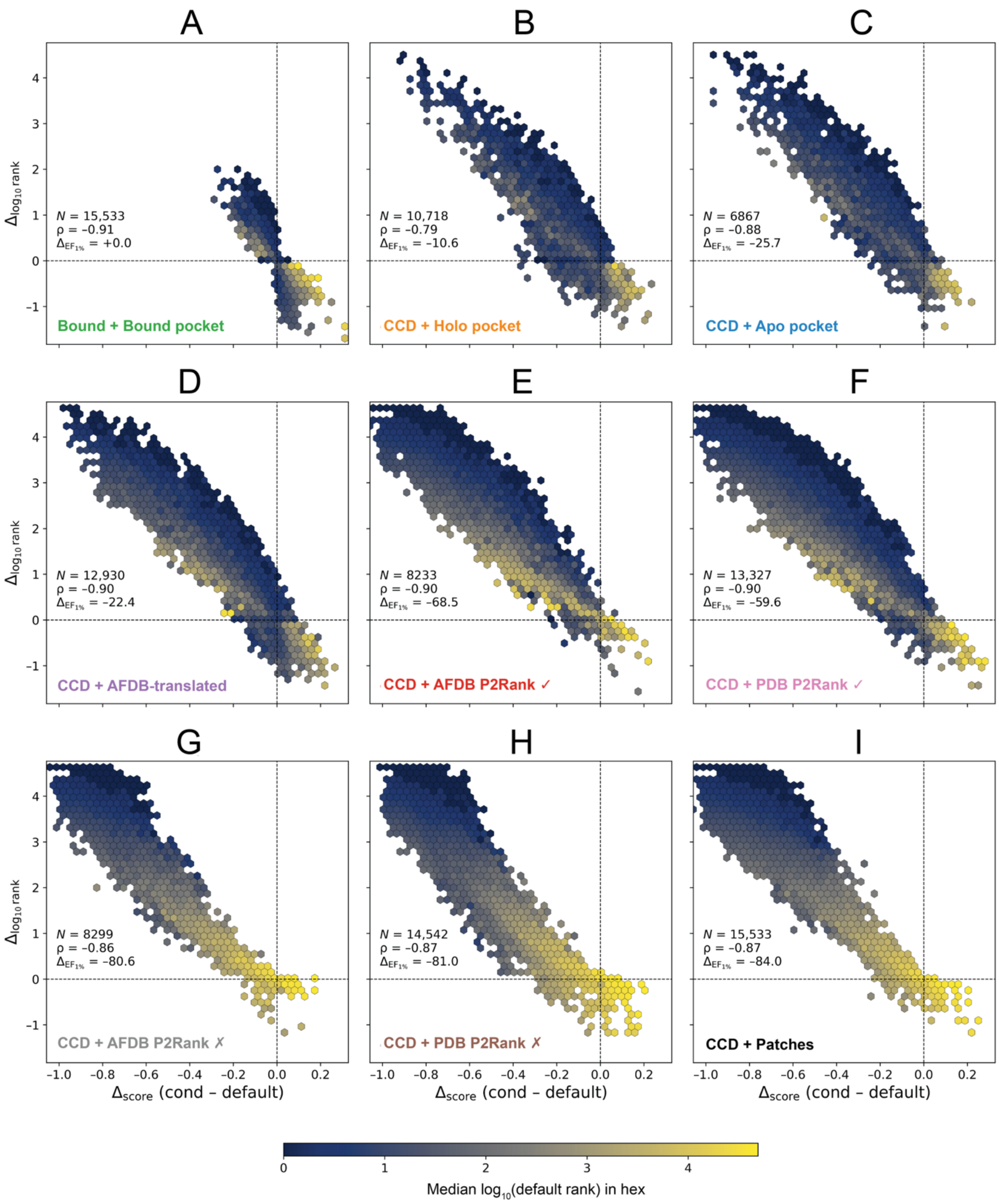


**Figure S4. Relationship between changes in ligand retrieval rank and pocket-ligand cosine similarity across screening conditions.** Hexbin plots showing the relationship between changes in the cosine similarity score of the crystal bound ligand ($\Delta_{score}$) and changes in its $\log_{10}$-transformed retrieval rank ($\Delta_{\log_{10} rank}$), relative to the default screening condition using a generic CCD ligand conformer and the experimental pocket. All comparisons were performed on the full pooled set of pockets ($L_0$), without applying any de-leakage filters. Negative $\Delta_{score}$ values indicate reduced pocket-ligand embedding similarity, and positive $\Delta_{\log_{10} \text{rank}}$ values indicate poorer ligand retrieval. **(A)** Bound ligand and bound pocket; **(B)** CCD ligand and alternative holo pocket; **(C)** CCD ligand and apo pocket; **(D)** CCD ligand and AFDB-translated pocket; **(E, F)** CCD ligand and correctly predicted P2Rank pockets from AFDB models and experimental structures, respectively; **(G, H)** CCD ligand and incorrectly predicted P2Rank pockets from AFDB models and experimental structures, respectively; **(I)** CCD ligand and random surface patches. Hexagons are coloured according to the median $\log_{10}$-transformed retrieval rank of the known ligand under the default condition. The number of evaluated pockets (*N*), Spearman's rank correlation coefficient (ρ) between score and rank changes, and percentage change in enrichment factor at 1% ($\Delta_{EF_{1\%}}$) relative to the default condition are reported for each panel. $\Delta_{EF_{1\%}}$ was calculated using the same subset of pockets for both conditions in each comparison and may therefore differ from values derived from the results reported in Table S6. Dashed lines indicate no change relative to the default condition.

| Condition | $L_0$ | $L_1$ | $L_2 < 0.7$ | $L_2 < 0.5$ | $L_2 < 0.3$ | $L_3 < 100$ | $L_3 < 25$ | $L_3 < 5$ | $L_3 = 0$ | $L_4 < 0.9$ | $L_4 < 0.7$ | $L_4 < 0.5$ | $L_4 < 0.3$ |
|---|---|---|---|---|---|---|---|---|---|---|---|---|---|
| CCD + Bound | 15,533 | 9323 | 2705 | 1908 | 958 | 625 | 537 | 423 | 284 | 241 | 223 | 136 | 9 |
| Bound + Bound | 15,640 | 9412 | 2728 | 1925 | 961 | 628 | 540 | 425 | 286 | 241 | 223 | 136 | 9 |
| CCD + Holo | 10,718 | 6531 | 1658 | 1125 | 525 | 383 | 336 | 272 | 191 | 172 | 162 | 105 | 8 |
| CCD + Apo | 6867 | 3458 | 567 | 403 | 194 | 125 | 111 | 75 | 51 | 40 | 37 | 14 | 2 |
| CCD + AFDB-translated | 12,930 | 7828 | 2235 | 1557 | 731 | 504 | 437 | 344 | 244 | 205 | 191 | 121 | 8 |
| CCD + PDB P2Rank ✓ | 13,327 | 7866 | 2313 | 1618 | 811 | 522 | 451 | 365 | 244 | 211 | 197 | 124 | 9 |
| CCD + AFDB P2Rank ✓ | 8233 | 4807 | 1402 | 999 | 412 | 292 | 258 | 219 | 164 | 139 | 131 | 92 | 7 |
| CCD + PDB P2Rank ✗ | 14,542 | 8846 | 2633 | 1851 | 937 | 609 | 524 | 412 | 275 | 236 | 218 | 134 | 7 |

| | | | | | | | | | | | | | |
|---|---|---|---|---|---|---|---|---|---|---|---|---|---|
| CCD + AFDB P2Rank ✗ | 8299 | 4800 | 1381 | 974 | 418 | 305 | 267 | 226 | 167 | 141 | 133 | 94 | 6 |
| CCD + Patches | 15,533 | 9323 | 2705 | 1908 | 958 | 625 | 537 | 423 | 284 | 241 | 223 | 136 | 9 |

**Table S5. Sample size throughout the de-leakage ladder.** Number of evaluated pockets at each de-leakage ladder level across the ten conditions tested. $L_0$ corresponds to the full set. In $L_1$ same PDB matches are removed. $L_2$ considers protein sequence identity, $L_3$ observed ligand frequency, and $L_4$ chemical (ECFP4) fingerprint similarity to the protein-ligand complexes in the BioLiP [30] subset used to fine-tune DrugCLIP.

| **Condition** | **$L_0$** | **$L_1$** | **$L_2 < 0.7$** | **$L_2 < 0.5$** | **$L_2 < 0.3$** | **$L_3 < 100$** | **$L_3 < 25$** | **$L_3 < 5$** | **$L_3 = 0$** | **$L_4 < 0.9$** | **$L_4 < 0.7$** | **$L_4 < 0.5$** | **$L_4 < 0.3$** |
|---|---|---|---|---|---|---|---|---|---|---|---|---|---|
| CCD + Bound | 84.8 | 77.0. | 74.7 | 75.1 | 72.6 | 65.3 | 64.6 | 63.8 | 59.9 | 56.8 | 56.5 | 58.1 | 55.6 |
| Bound + Bound | 84.8 | 77.0 | 75.0 | 75.4 | 73.2 | 65.6 | 64.8 | 64.2 | 61.2 | 57.7 | 56.5 | 57.4 | 55.6 |
| CCD + Holo | 74.3 | 67.6 | 63.1 | 63.2 | 57.9 | 52.2 | 50.0 | 53.3 | 53.4 | 50 | 50 | 54.3 | 37.5 |

| | | | | | | | | | | | | | |
|---|---|---|---|---|---|---|---|---|---|---|---|---|---|
| CCD + Apo | 61.2 | 51 | 39.9 | 42.7 | 37.1 | 32 | 32.4 | 36.0 | 33.3 | 37.5 | 40.5 | 50.0 | 0.0 |
| CCD + AFDB-translated | 61.8 | 56.2 | 57.1 | 58.2 | 53.4 | 44.4 | 43.5 | 47.1 | 45.1 | 41.0 | 39.8 | 40.5 | 50 |
| CCD + PDB P2Rank ✓ | 27.1 | 23.2 | 17.3 | 16.7 | 15.9 | 13.0 | 12.0 | 11.2 | 12.7 | 11.8 | 10.7 | 9.7 | 0.0 |
| CCD + AFDB P2Rank ✓ | 16.8 | 16.4 | 13.3 | 13.2 | 12.9 | 11.3 | 12.8 | 11.0 | 8.5 | 7.9 | 8.4 | 5.4 | 0.0 |
| CCD + PDB P2Rank ✗ | 3.4 | 3.7 | 2.6 | 3.2 | 3.5 | 4.9 | 4.4 | 3.9 | 2.2 | 2.5 | 2.8 | 1.5 | 0.0 |
| CCD + AFDB P2Rank ✗ | 2.7 | 3.0 | 2.8 | 2.6 | 3.3 | 3.9 | 3.7 | 2.7 | 2.4 | 2.1 | 1.5 | 1.1 | 16.7 |
| CCD + Patches | 0.8 | 0.8 | 1.0 | 1.3 | 1.5 | 2.2 | 2.6 | 3.1 | 1.1 | 1.2 | 1.3 | 1.5 | 0.0 |

**Table S6. $EF_{1\%}$ throughout the de-leakage ladder.** DrugCLIP's performance ($EF_{1\%}$) at each de-leakage ladder level across the ten conditions tested.

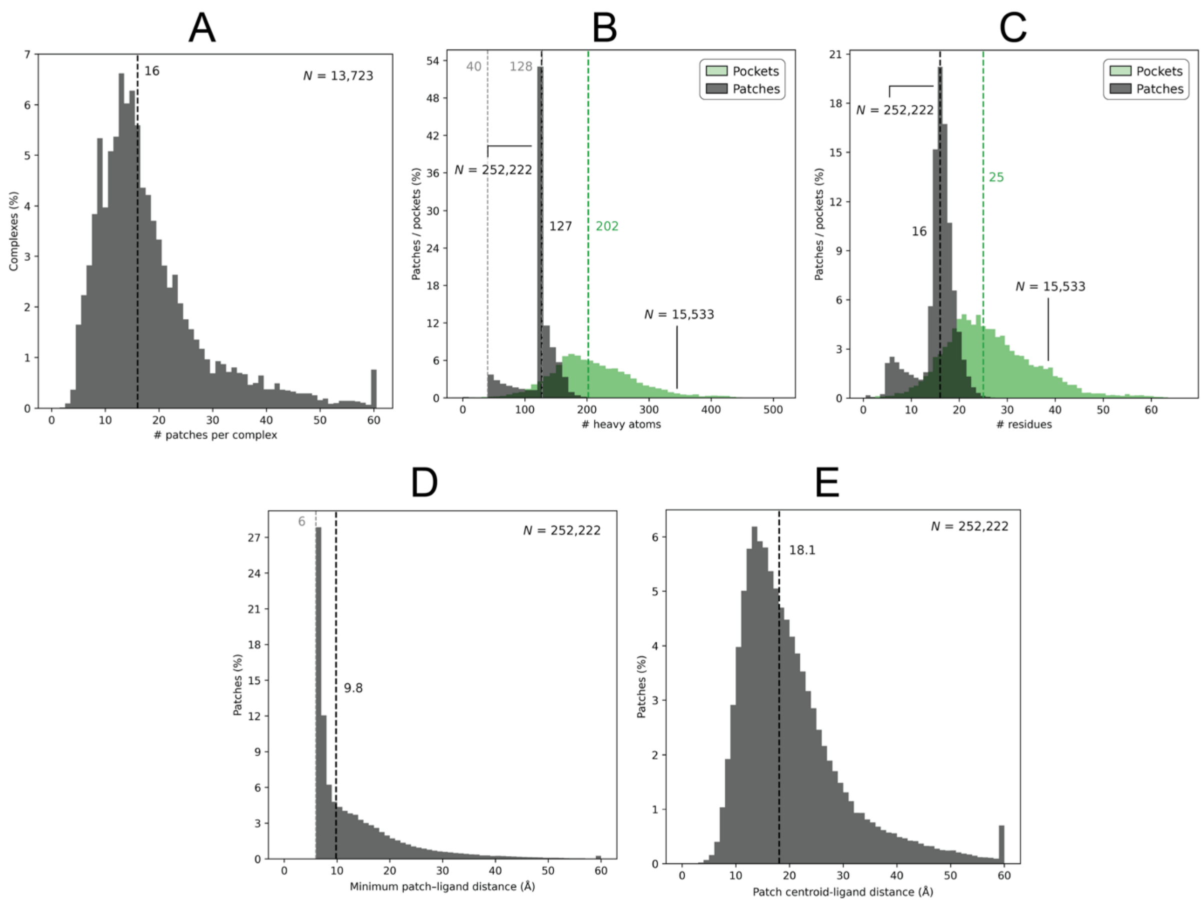


**Figure S5. Negative control random surface patches characterisation.** Statistics are pooled across COACH420 [31], HOLO4K [32], scPDB [33] and PDBBind [34] for the cross-data set deduplicated screening set comprising 13,723 complexes, 15,533 experimental ligand-bound pockets and 252,222 patches. **(A)** Distribution of the number of random surface patches generated per protein-ligand complex or structure; **(B)** Distribution of the number of heavy atoms in random patches (grey) and experimental ligand-binding pockets (green); **(C)** Distribution of the number of amino acid residues in random patches and experimental ligand-binding pockets; **(D)** Distribution of the minimum distance between any random patch heavy atom and the corresponding bound ligand heavy atoms in the complex; **(E)** Distribution of the distance between random patch centroids and the corresponding ligand centroids. Grey dashed vertical lines mark the thresholds employed in the patch generation algorithm, i.e., 6 Å, 128 and 40 atoms, while coloured ones indicate median values, and *N* denotes the number of observations for each set.

| | |
|---|---|
| # complexes (total) | 13,723 |
| # pockets (total) | 15,533 |
| # patches (total) | 252,222 |
| # patches per complex | 16 – IQR = [11-22] |
| # heavy atoms per patch | 127 – IQR = [124-135] |
| # residues per patch | 16 – IQR = [15-18] |
| Patches < 40 atoms (%) | 0.3 |
| Patches > 128 atoms (%) | 30.5 |
| # patch residues within 6 Å of ligand | 0 |
| Patches ≤ 8 Å from ligand (%) | 39.9 |
| Patch centroid-ligand distance (Å) | 18.1 – IQR = [13.6-24.5] |
| Minimum patch-ligand distance (Å) | 6.0 |
| # heavy atoms per pocket | 202– IQR = [164-250] |
| # residues per pocket | 25 – IQR = [20-32] |

**Table S7. Negative control random surface patches summary statistics.** Statistics are pooled across COACH420, HOLO4K, scPDB and PDBBind for the cross-dataset deduplicated screening set (13,723 complexes; 15,533 ligand pockets; 252,222 patches). Continuous variables are given as median – [interquartile range (IQR)]. Pockets are the experimental ligand binding sites, i.e., all amino acid residues with at least one atom within 6 Å of a bound ligand heavy atom, as per the DrugCLIP pocket definition.
